# Structure-Guided Identification of Critical Residues in the Vacuolar Na^+^,K^+^/H^+^ Antiporter NHX1 from *Arabidopsis thaliana*

**DOI:** 10.1101/2022.05.18.492413

**Authors:** Belen Rombola-Caldentey, Imelda Mendoza, Francisco J. Quintero, José M. Pardo

**Author notes:** Author for correspondence: Jose M. Pardo. **Author Contributions:** F.J.Q. and J.M.P designed and supervised the research; B.R.C. and I.M. performed experiments and analyses; all authors analyzed and discussed the data; B.R.C. and J.M.P wrote the manuscript; J.M.P. obtained the funding. All authors have read and agreed to the published version of the manuscript.

## Abstract

Cation/Proton Antiporters (CPA) acting in all biological membranes help regulate the volume and pH of cells and of intracellular organelles. A key issue with these proteins is their structure-function relationships since they present intrinsic regulatory features that rely on structural determinants, including pH-sensitivity and the stoichiometry of ion exchange. Crystal structures are only available for prokaryotic CPA, whereas the eukaryotic ones have been modeled using the former as templates. Here we show an updated and improved structural model of the tonoplast-localized K^+^,Na^+^/H^+^ antiporter NHX1 of Arabidopsis as a representative of the vacuolar NHX family that is key to the accumulation of K^+^ into plant vacuoles. Conserved residues judged as functionally important were mutated and the resulting protein variants were tested for activity in the yeast *Saccharomyces cerevisiae*. Results indicate that residue N184 in the ND-motif characteristic of CPA1 could be replaced by the DD-motif of CPA2 family members with minimal consequences on activity, yet this residue may help to regulate the optimal pH range of the exchanger. Attempts to alter the electroneutrality of AtNHX1 by different combinations of amino acid replacements at N184, R353 and R390 residues resulted in inactive or partly active proteins with differential ability to control the vacuolar pH of the yeast.

## INTRODUCTION

Cation/Proton Antiporters (CPAs) play a major role in regulating the volume and pH of cells and of intracellular organelles. These antiporters mediate the exchange of monovalent cations Na^+^ and K^+^ with one or two H^+^ across the membrane with different stoichiometry. Consequently, CPAs are divided into two main groups, CPA1 and CPA2, whose members often differ in their ion selectivity and electrogenicity. A key issue with these proteins is their structure-function relationships since many CPA proteins present intrinsic regulatory features that are intimately connected to structural determinants. High-resolution structures are only available for CPA proteins of prokaryotic origin, and they have served as template to model the eukaryotic counterparts. All CPA proteins are thought to share a similar transmembrane topology at the active center, known as the Nha-fold (Masrati et al. 2018), in which two short, partly-unwound helices in transmembranes 4 and 11 create two partial positive (TM4c and TM11p) and two partial negative (TM4p and TM11c) dipoles facing each other (Călinescu et al. 2017; Hunte et al. 2005).

Most CPA1 proteins contain a conserved asparagine-aspartate pair at their Nha-fold (ND-motif) in TM5 whereas CPA2 members present an aspartate-aspartate pair (DD-motif). This dichotomy in the CPA1/CPA2 division has often been correlated with electrogenicity so that the DD-motif was thought to allow the simultaneous translocation of two H^+^. However, this classical view has been challenged recently. Extensive phylogenetic comparison together with structure-guided mutagenesis has indicated the CPA1/CPA2 division only partially correlates with electrogenicity, and that this property is not due to this DD-motif as previously thought (Masrati et al. 2018). A rationally designed triple mutant, only one of which was a change from the DD-motif to the ND-motif, successfully converted the electrogenic EcNhaA to be electroneutral (Masrati et al. 2018). The electrogenicity of the CPA2 family member NapA from *Thermus thermophilus* relied on the presence of a conserved Lys residue in TM10, which in electroneutral CPA1 members is exchanged for an Arg (Uzdavinys et al. 2017). Further, HsNHA2 is an electroneutral protein despite of the DD-motif in the active center and has an Arg residue (R432) in the homologous position to K305 of the electrogenic NapA from *Thermus thermophilus*. Mutation K305R changed NapA to electroneutrality, but the converse mutation R432K in HsNHA2 failed to trigger electrogenic transport in (Uzdavinys et al. 2017). The lack of consistent outcomes from structure-function studies keeps the basis for electrogenic transport in CPA exchangers uncertain.

To date, no crystallographic structure has been determined for any eukaryotic NHE/NHX protein of the CPA1 family. Nonetheless, different topological models have been described for the Arabidopsis K^+^,Na^+^/H^+^ exchanger NHX1, with contradictory results (Sato and Sakaguchi 2005; Yamaguchi et al. 2003). On the one hand, Yamaguchi et al (2003) proposed a model in which the N-terminus of the protein is cytosolic while the C-terminus is facing the vacuolar lumen; according to this model, the hydrophobic domain of AtNHX1 presented nine transmembrane domains and three hydrophobic regions that appeared to be membrane-associated. By contrast, Sato and Sakaguyi (2005) proposed a model in which both N- and C-termini were cytosolic, and the pore domain was arranged as 12 transmembrane segments with an intramembranous loop, i.e. not crossing the membrane entirely. Although the latter model is more consistent with previous studies with other proteins of the CPA superfamily, the study of Sato and Sakaguyi (2005) may be questioned by the use of fragments of AtNHX1 rather than the full-length protein. The main controversy in these reports is the orientation of the C-terminal tail. However, the presence of phosphorylated residues within the C-terminal tail of the mammalian HsNHE1, the fungal protein ScNHX1, and in the Arabidopsis NHX1/2 protein has been described (Albuquerque et al. 2008; Gruhler et al. 2005; Whiteman et al. 2008), implying that their C-termini must be cytosolic as no intravacuolar protein phosphorylation has been described so far.

Using prediction tools and *in silico* modeling, the topology and 3D models of several plant proteins of the CPA family have been generated based on known atomic structures of prokaryotic Na^+^/H^+^ antiporters that are phylogenetically related. Tridimensional structures have been generated for the tree *Populus euphratica* PeNHX3 using as template the structure of EcNhaA from *E. coli* (Wang et al. 2014) and for CHX17 using the TtNapA structure of *Thermus thermophilus* as the template (Czerny et al. 2016). Moreover, the Arabidopsis AtNHX6 structure was modeled using the MjNhaP of *Methanococcus jannaschii* as the template (Ashnest et al. 2015). However, the confidence of these structures was not assessed by experimental approaches to corroborate the structures proposed. In this study, a model structure for AtNHX1 was generated to better understand the structure-function relationships of the protein and a mutagenic analysis was carried out with conserved residues presumed to be important for protein activity. Here we show that the ternary model of AtNHX1 conserves the structural features characteristic of microbial and mammalian members of the CPA superfamily. Several conserved amino acids that are essential for the activity of AtNHX1 have been identified at the active site and the Nha-fold. These residues are the T156 and D157 of the TD-motif in TM4, D185 of the ND-motif in TM5, and arginines R353 and R390 in TM10 and TM11, respectively. Residue N184 at the ND-motif that is highly conserved in electroneutral antiporters of the CPA1 family and, surprisingly, it is not essential for the activity of AtNHX1, yet it may help regulate the optimal pH range for protein activity. Last, in agreement with the proposed function *in planta*, we show that AtNHX1 provides a H^+^ shunt at the tonoplast that regulates vacuolar pH in yeasts cells.

## RESULTS

### Topological model

The AtNHX1 protein sequence was used as query in the SwissModel web-server dedicated to homology modeling of protein structures (Bienert et al. 2017; Waterhouse et al. 2018). This software generated a list of proteins with known structure that were the best fits in terms of sequence conservation with the query sequence. Top-ranked templates were the archaea proteins NhaP of *Pyrococcus abysinnia* (Wöhlert et al. 2014), NhaP1 of *Methanocaldococcus jannichii* (Goswami et al. 2011), and NapA of *Thermus thermophilus* (Lee et al. 2013b). All these archaeal proteins had better scores than the archetypical bacterial protein NhaA of *E. coli* (Lee et al. 2014) (Table S1). It should be noted that the AtNHX1 sequence in these alignments included mainly the N-terminal hydrophobic domain of the transporter, with a coverage ranging 58-71% of the protein length, while the C-terminal was not aligned in any case due to the lack of sequence similarities between the eukaryotic protein and the prokaryotic proteins. In order to refine the results, the protein alignment was repeated using as the query only amino acids 1-435 of AtNHX1 constituting the pore domain of the protein. The results obtained, although with better scores, did not depart from the previous ones (Table S2). To decide which template to select for further analyses, the following basic parameters were established: the structure had been obtained by X-ray, the resolution should at least be 2.9 Å, and in case of duplication of template structures the one with the highest score. The list was then reduced to four templates for further analyses: 4cza.1 (PaNhaP from *Pyrococcus abyssii*), 4czb.1 (MjNhaP1 from *Methanocaldococcus jannaschii*), 4bwz.1.A (TtNapA from *Thermos thermophillus*); and Aau5.1.A (EcNhaA, from *Escherichia coli*).

To obtain a consensus topological model for the pore domain of AtNHX1, the template protein sequences selected above where aligned pairwise with AtNHX1 using ClustalW software. Each of these alignments was used to assign the boundaries of the transmembranes of AtNHX1 based in the protein topology in the crystal structure of the templates. Finally, the sequences of AtNHX1 with the assigned secondary structures of each alignment were compared (Figure S1). The structure of the NhaP-type proteins comprised thirteen complete TM segments instead of the twelve in EcNhaA. From these alignments we concluded that the EcNhaA template was not appropriate to model AtNHX1. Firstly, large segments in the N-terminal and C-terminal portions of the AtNHX1 sequence were not covered by the alignment, while they were present in the alignment of the phylogenetically closer structures of PaNhaP and MjNhaP1. As a result, the alignment of AtNHX1 with EcNhaA did not allow the location of the complete hydrophobic domain of the protein. However, the other three templates not only allowed the alignment of the complete sequence, but also the predicted segments in each case had a similar arrangement of transmembranes regardless of the template used. Consequently, the final topological model was generated based in the alignment of AtNHX1 with PaNhaP, MjNhaP1 and TtNapA templates (Figure 1A). The proposed topology of AtNHX1 consist of two well defined domains: a hydrophobic core that includes the largest portion of the protein (from residues A28 to T434) that corresponds to the pore domain involved in ion transport, and a hydrophilic and cytosolic C-terminal tail (from amino acids K435 to A538). In our model, both N- and C-termini are cytosolic and the complete C-terminal domain downstream the pore domain is in the cytosol. The N-terminal portion of the pore domain consist of two intramembranous (IM) semihelices (IM1.a and IM1.b) followed by eleven TM segments with an antiparallel topology, so that the odd TM segments (TM3, TM5, TM7, TM9 and TM11) are oriented from inside (cytosol) to outside (vacuole), while the even TM segments (TM2, TM4, TM6, TM8, TM10 and TM12) from outside to inside. These TM segments are connected by short loops of varied length, being the shorter of 4 amino acids (between TM4 and TM5), and the longest of 18 (between IM1.b and TM2). The C-terminal tail is the most variable region among plant NHX proteins and may integrate the signals for protein regulation, such as protein phosphorylation and binding of calmodulin-like proteins (Pabuayon et al. 2021; Whiteman et al. 2008; Yamaguchi et al. 2003).

**Figure 1.**
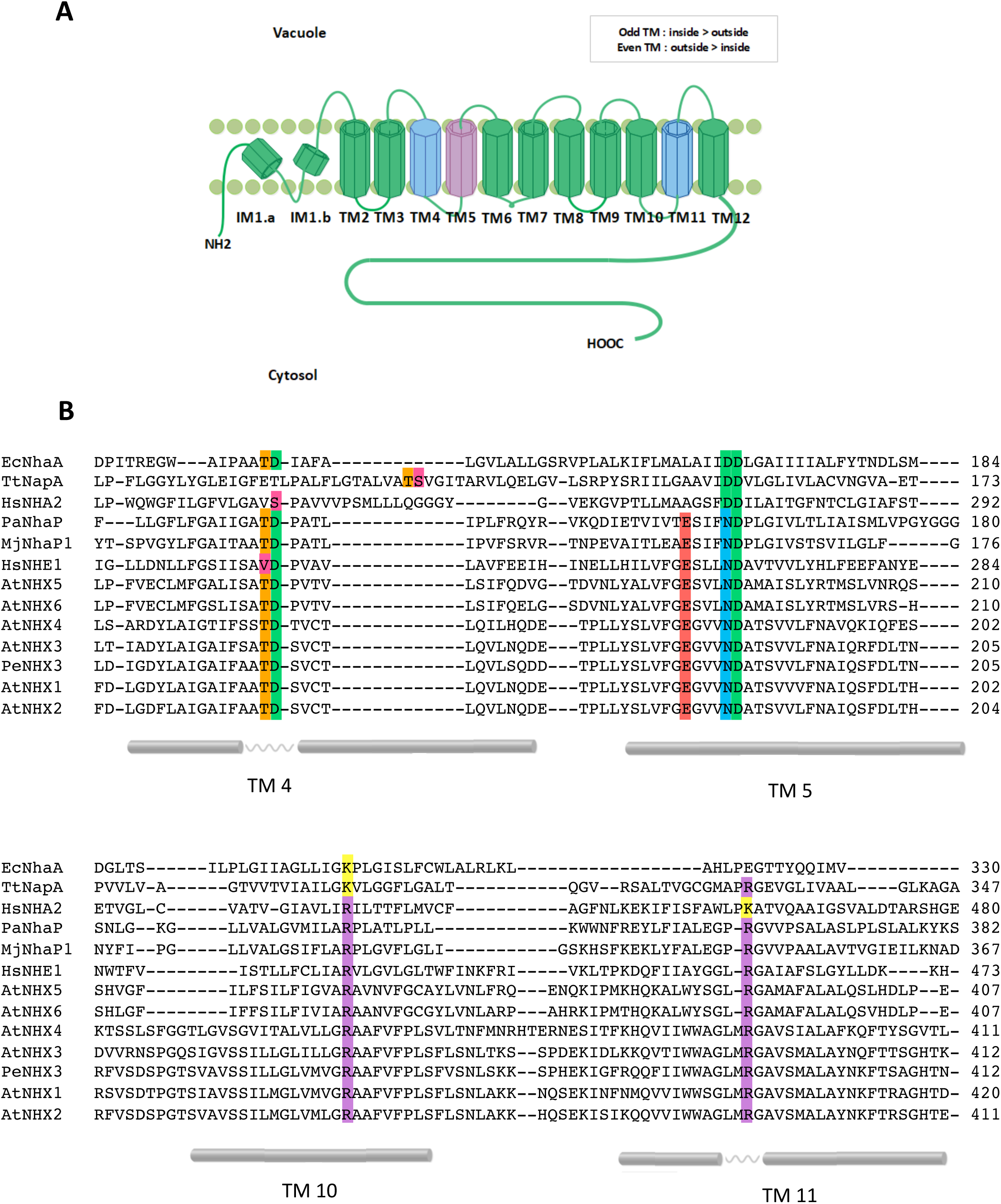
Topological model of AtNHX1 based in the alignment of AtNHX1 sequence with prokaryotic templates. **(A)** Schematic representation of AtNHX1 topology based on sequence alignments to templates PaNhaP, MjPhaP1 and TtNapA (see Supplemental Figure 1). AtNHX1 consist of two domains: a hydrophobic catalytic core and a C-terminus hydrophilic tail. The catalytic core is formed by two intramembranous (IM) semi-helices followed by eleven TM segments with antiparallel orientation. Both the N- and C-termini are cytosolic. **(B)** Sequence alignment of CPA1 and CPA2 proteins cited here. The most important conserved amino acids in the hydrophobic core and the TM segments where they are located are highlighted. TM numeration is according to the topological model of AtNHX1 shown in panel A.

### Tertiary structure model of AtNHX1

Using the modeling data retrieved from the SwissModel server and visualized with the Pymol software, the 3D structure model of the complete pore domain of AtNHX1 was generated (Figure 2). Among the SwissModel results, the two templates with the highest scores corresponded with two phylogenetically related archaea proteins PaNhaP and MjNhaP1 (Supplemental Tables S1 and S2), which have been shown to present similar structure and function (Călinescu et al. 2016; Călinescu et al. 2014). On the other hand, the TtNapA protein, despite being an electrogenic exchanger, is structurally more similar to PaNhap and MjNhaP1 than to EcNhaA (Hunte et al. 2005). For this reason, the AtNHX1 model generated using TtNapA as template was more similar to the ones obtained from PaNhaP and MjNhap1 as models compared to the EcNhaA template. The experimentally determined structures of PaNhaP and MjNhaP1 were aligned to each other, as well as the modeled structure of AtNHX1 for each template (Figure 2C). The differences observed between structures were negligible. We selected PaNhaP for further analyses because, unlike MjNhaP1, this protein is able to bind Ti^+^ (besides Na^+^ and Li^+^), whose ion radius (1.5 Ȧ) is similar to that of K^+^ (1.44Ȧ) and larger than that of Na^+^ (1.12 Ȧ) (Padan and Michel 2015). AtNHX1 has low Na^+^-K^+^ discrimination (Hernandez et al. 2009).

**Figure 2.**
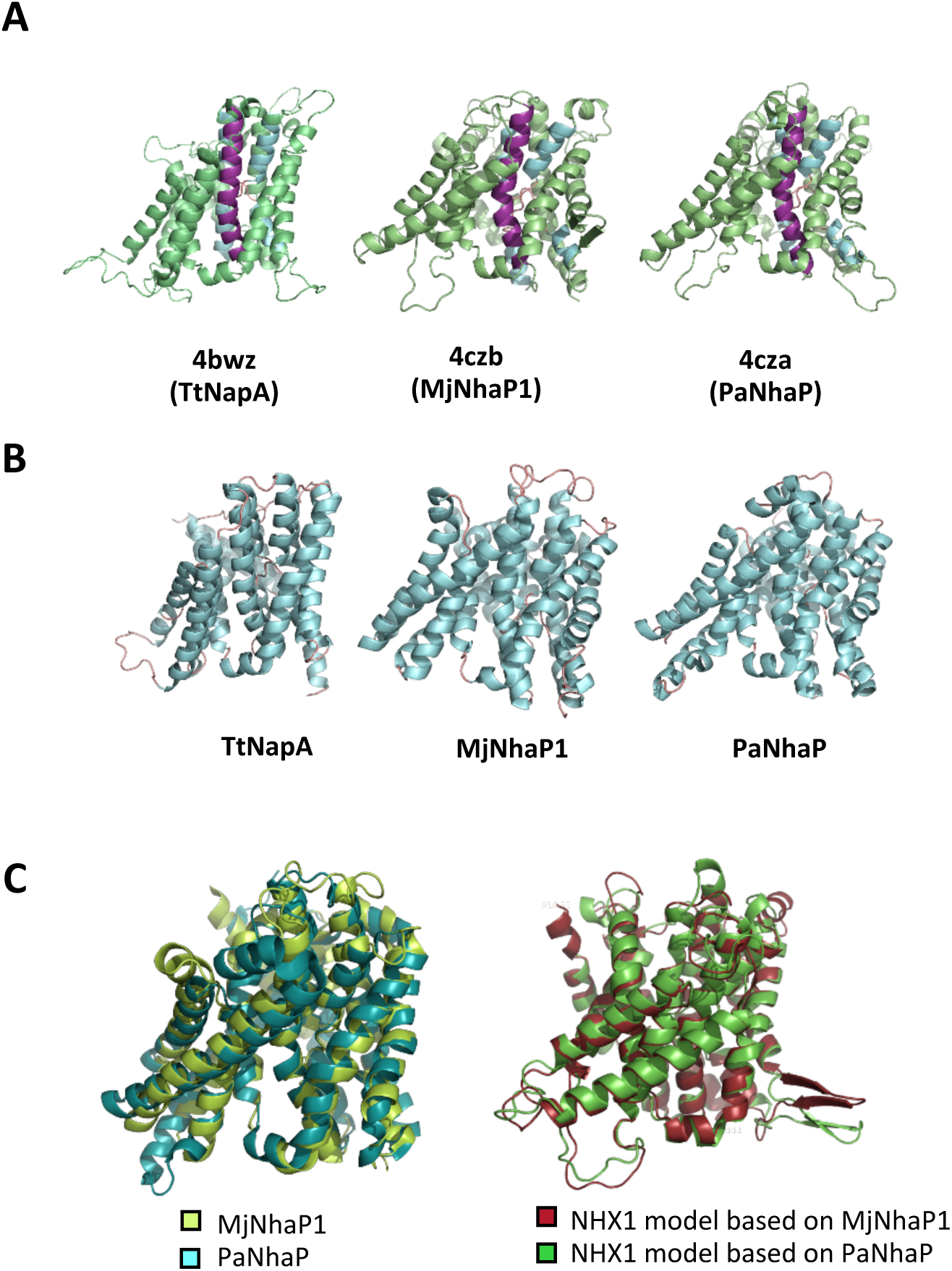
Modelization of AtNHX1 structure based on crystalized prokaryotic templates. **(A)** Tridimensional structure representation of AtNHX1 models produced by SwissModel using the structural templates indicated. TM5 is highlighted in purple; TM4 and TM11, in blue. **(B)** Crystal structure of the prokaryotic template proteins. **(C)** Structure overlay with Pymol software of the archea proteins MjNhaP1 and PaNhaP (left), and of the AtNHX1 models based in these template structures.

### Nha-fold in the active center of AtNHX1

The structural assembly of the active center for the EcNhaA protein, the first CPA family member to be crystallized, is known as the Nha-fold (Hunte et al. 2005; Padan et al. 2009). In this non-canonical conformation of TM helices, the helical secondary structure of TM4 and TM11 of EcNhaA is interrupted by extended chains in the middle of the membrane, leaving instead two short helices in each TM flanking the centrally located extended chains (Hunte et al. 2005; Lee et al. 2014). The interrupted helices of TM4 and TM11 comprising the Nha-fold cross each other at the extended chains in the proximities of the active site, which is located in the TM5 of EcNhaA (Supplemental Figure S2). As a result, two short helices oriented either toward the cytoplasm (c) or toward the periplasm (p) create two partial positive (TM4c and TM11p) and two partial negative (TM4p and TM11c) dipoles facing each other (Călinescu et al. 2017; Hunte et al. 2005). These charges are compensated by Asp133 in TM4 and Lys300 in TM10, respectively (Lee et al. 2014), creating a balanced electrostatic environment in the middle of the membrane at the ion binding site. This fold it has been confirmed for all the crystallized CPA superfamily members (Hunte et al. 2005; Lee et al. 2013b; Lee et al. 2014; Paulino et al. 2014; Wöhlert et al. 2014) and incorporated into modeled structures (Landau et al. 2007; Schushan et al. 2010; Wang et al. 2014). Figure 3 illustrates the modeled TM segments forming the Nha-fold of AtNHX1 using PaNhaP as template, whereas Supplemental Figure S3 shows the models based on templates MjNhaP1 and TtNapA.

**Figure 3.**
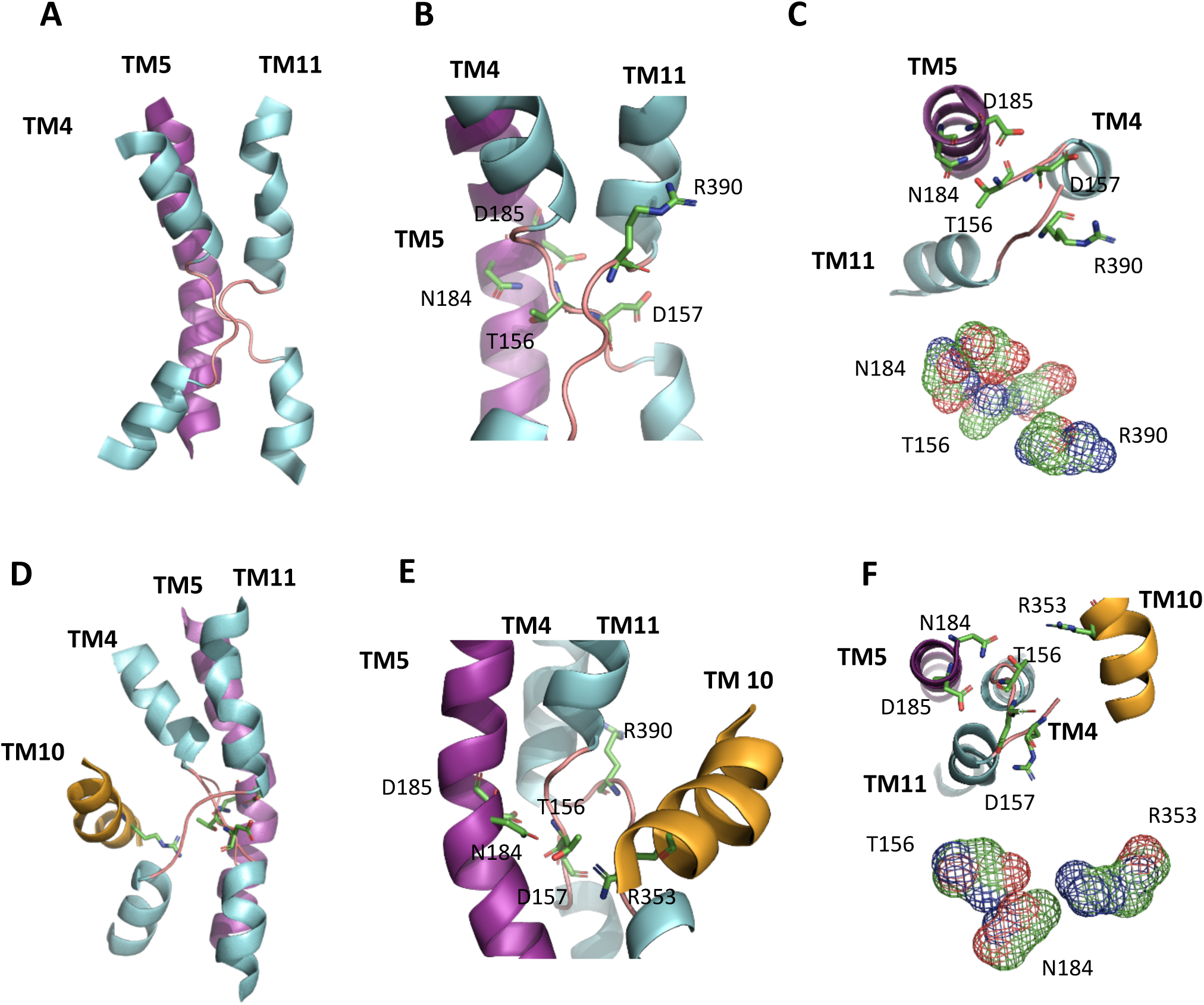
Nha-fold in the active center of AtNHX1. **(A)** Representation of TM4, 5 and 11 in the AtNHX1 structure showing the Nha-fold in the active center of the protein, modeled with PaNhaP as template. **(B),** Close-up view of the cross-over of the extended chains in TM4 and TM11 indicating the highly conserved residues T156 and D157 in TM4, N184 and D185 in TM5, and R390 in TM11. **(C)** A top-view (inside to outside) to show how D157 and R390 side chains compensate the dipoles generated by the crossing of TM4 and TM11. Nearby, T156 and N184 side chains are close enough to interact. **(D)** Representation of the active center with the Nha-fold and TM10. **(E)** Closer view of the Nha-fold and the localization of R353 in TM10 relative to the active center**. (F)** A top-view (inside to outside) to show the interaction of N184, T156 and R353 side chains.

### Conserved residues in the pore domain

Alignment of the primary structure of representative proteins of the CPA superfamily allowed the identification of highly conserved amino acids in the active center comprising the Nha-fold (TM4, TM5 and TM11) and the adjacent TM10 (Figure 1B and Figure 3). The aspartic acid at position 185 (D185) and the asparagine at 184 (N184) in TM5 comprise the active site of the protein. This ND-motif is highly conserved among electroneutral CPA1 family members, as described for MjNhaP1 (N160-D161, (Paulino et al. 2014), PaNhaP (N158-D159, (Paulino et al. 2014), HsNHE1 (N266-D267, (Landau et al. 2007) and all six members of the Arabidopsis NHX family (Figure 1B). In the electrogenic members of the CPA2 family, e.g. EcNhaA and TtNapA, this ND-motif is exchanged for a DD-motif (D163-D164 and D200-D201, respectively) (Hunte et al. 2005; Lee et al. 2013b; Padan et al. 2009) (Figure 1B). These results are consistent with the placement of AtNHX1 in the CPA1 clade and electroneutral exchange activity (Gradogna et al. 2021; Venema et al. 2002).

Two highly conserved, positively charged amino acids, R353 in TM10 and R390 in TM11, were detected in AtNHX1 (Figure 1B and Figure 3). Other electroneutral proteins of the CPA1 family also present conserved basic residues at equivalent positions, e.g. R425 and R458 in HsNHE1, and R432 and K460 in HsNHA2 (Landau et al. 2007; Schushan et al. 2010; Uzdavinys et al. 2017). In MjNhaP1 and PaNhaP, the first arginine is conserved in TM11 (equivalent to TM10 of AtNHX1) as residues R320 and R337, respectively. The crystal structures have shown that this conserved arginine interacts with a glutamate in TM6 (E156 in MjNhaP1 and E154 in PaNhaP; equivalent to E180 in TM5 of AtNHX1) that is also conserved in the CPA1 antiporters (Figure 1B). In the models obtained for AtNHX1 based on templates PaNhaP and MjNhaP1, R353 seems to interact with E180 (Figure 4B,C). Moreover, E180 interacts with N184 in the three AtNHX1 models shown in Figure 4. The second arginine, R390 in TM11 of AtNHX1, is also conserved (Figure 1B), but little is known about the function of this conserved residue. In HsNHA2 this residue is exchanged for a Lys.

**Figure 4.**
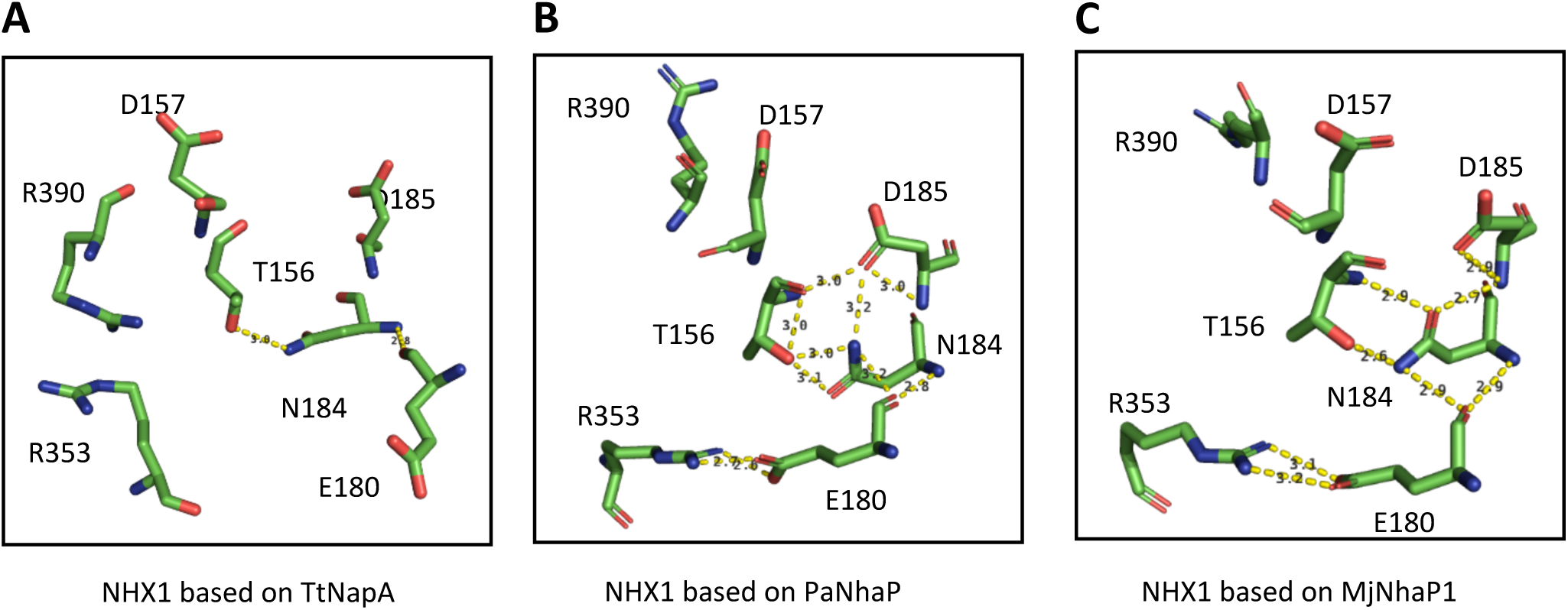
Predicted amino acid interactions in the active site. Representation of the side chains of the amino acids proposed to be affecting the activity or structure of the AtNHX1 protein according to the model obtained from the **(A)** TtNapA, **(B)** PaNhaP and **(C)** MjNhaP1 templates. Yellow dashes indicate the possible interactions based in the distance between residues.

Conserved threonine and aspartate residues in TM4 form a TD-motif in the unwound chain, which is also conserved in EcNhaA (T132-D133) (Galili et al. 2002; Maes et al. 2012), and in the archea proteins MjNhaP1 and PaNhaP. In both cases, bacterial and archea proteins, this TD-motif has been described to take part in ion coordination (Paulino et al. 2014; Wöhlert et al. 2014). Moreover, T129 and T131 of PaNhaP and MjNhaP1 interact by their side chain with residues N158 and N160, respectively. These asparagines do not participate in the coordination of the substrate ion but they control the access to the ion-binding site (Paulino et al. 2014; Wöhlert et al. 2014). In the mammalian HsNHE1 and HsNHA2 and in prokaryotic TtNapA this motif is not conserved (Landau et al. 2007; Lee et al. 2013b; Schushan et al. 2010). In the modeled structure of PeNHX3 no conserved TD-motif was described, and instead the presence of a non-charged Tyr in position 149 was proposed to be substitute for the Thr residue in the TD-motif (Wang et al. 2014). However, the alignment of the CPA proteins shown in Figure 1B indicates that the TD-motif is conserved in PeNHX3. In AtNHX1 this motif is conserved as well (T156-D157), and according to all the models obtained for AtNHX1, there seems to be an interaction of T156 with N184, as described previously for the archaea NhaP proteins (Figure 4).

### In silico validation of the AtNHX1 structural model

To evaluate the quality of the structural model generated for AtNHX1, several in silico analyses were used (Kondapalli et al. 2013; Landau et al. 2005; Schushan et al. 2010; Wang et al. 2014). These analyses include generic characteristics commonly displayed in membrane proteins, such as the distribution of positively charged residues and the pattern of evolutionary conservation (Bowie 2005; Fleishman and Ben-Tal 2006).

The topology of most of the intrinsic membrane proteins is such that intracellular positions are enriched with positively charged residues (Arg and Lys) compared to extracellular regions (Wallin and von Heijne 1998). This is known as the “positive-inside rule” and has been used to evaluate the quality of structural models of membrane proteins, including the structures of the antiporters EcNhaA, HsNHE1, HsNHA2 or PeNHX3 (Hunte et al. 2005; Landau et al. 2007; Schushan et al. 2010; Wang et al. 2014). Analysis of charge distribution in the AtNHX1 three-dimensional structure modeled with templates TtNapA, MjNhaP1 and PaNhaP confirmed this rule (Figure 5A). When the PaNhaP1-derived model was taken into consideration, out of 25 Arg and Lys residues in the AtNHX1 protein, 14 residues are in the cytosolic side of the membrane, 9 in the outside (vacuolar lumen), and 2 are intramembranous (Figure 5A). As expected for an intrinsic membrane protein, the predicted AtNHX1 structure showed that the polar residues were clustered either in the inner structures of the protein or at the extramembranous loops, while maximizing the exposure of hydrophobic residues to the membrane lipids (Figure 5B).

**Figure 5.**
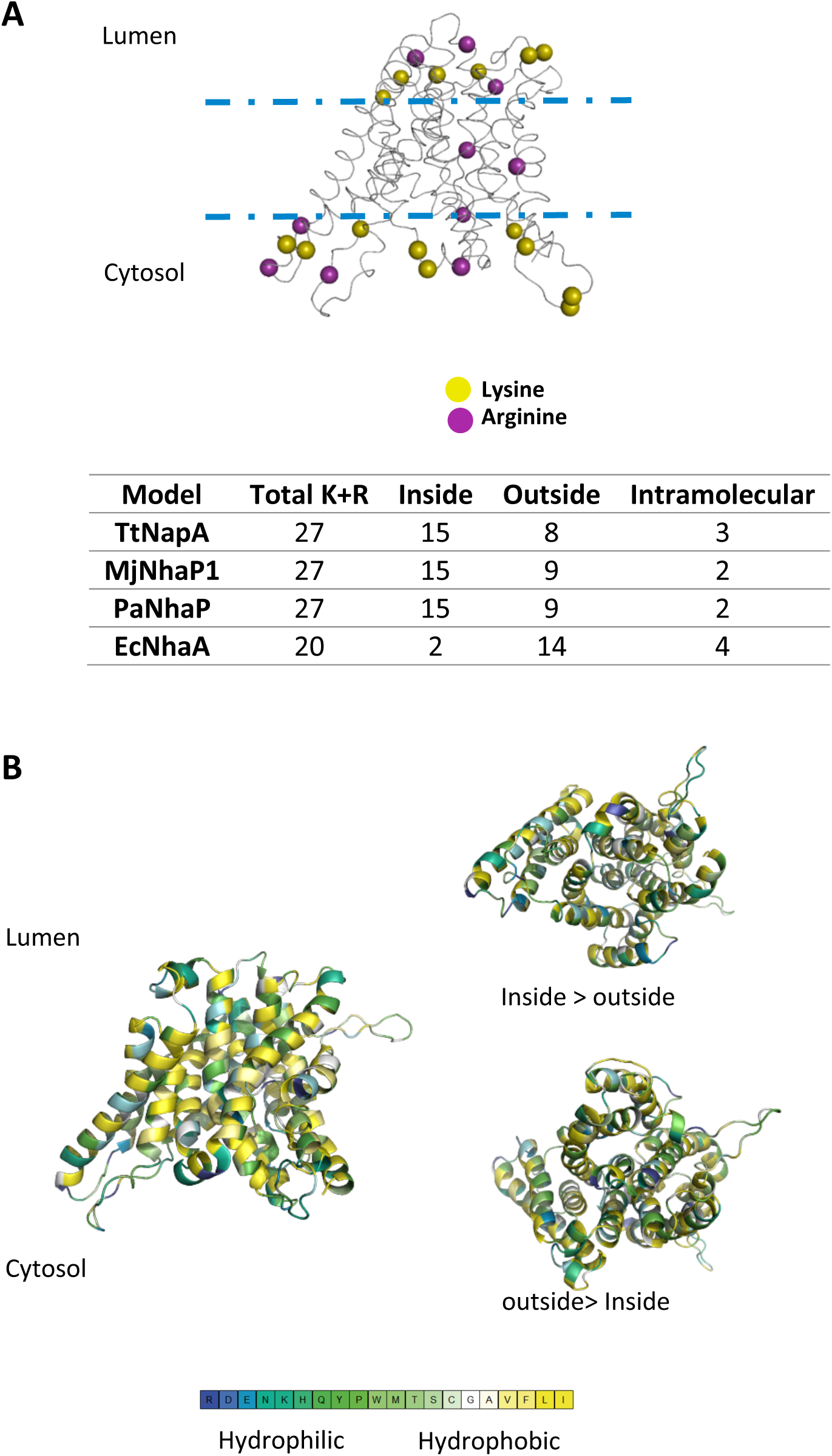
Validation of the modeled AtNHX1 structure. The AtNHX1 model represented corresponds to the PaNhaP template**. (A)** The Cα of lysines (yellow) and arginines (purple) are highlighted in the structure to determine their distribution in the protein. Except for the model based on EcNhaA, other modeled structures of AtNHX1 are in accordance with the ‘positive-inside’ rule. **(B)** Distribution of the amino acids according to their hydrophobicity across the AtNHX1 structure based on the PaNhaP template. Amino acids are colored based on the hydrophobicity scale of Kessel and Ben-Tal (2002).

Evolutionary conservation of amino acids has been previously used to assist structure modeling. Ion transporters are subjected to evolutionary pressure mainly restricting amino acid changes in regions that take part in ion binding or translocation, but also of inter-helical interfaces, crucial for stabilizing the architecture of the helix bundle. For that reason, it is expected that the protein core would be conserved whereas residues that face the lipids or are in the cytosolic and luminal loops were more variable (Fleishman and Ben-Tal 2002). The evolutionary conservation for AtNHX1 was calculated using the ConSurf server (http://consurf.tau.ac.il/) (Landau et al. 2005). To generate a conservation model, three different approaches with increasing levels of stringency were used. In brief, AtNHX1 sequence was used as a query in the UNIPROT database (Bairoch et al. 2005) using PSI-BLAST (Altschul et al. 1997) to collect homologous sequences. Redundant (>95% sequence identity) or with low coverage of the protein (< 60% identity) as well as fragmented sequences were discarded. The resulting 216 sequences were aligned using MUSCLE (Edgar 2004) with default parameters, and the final Multi-Sequence Alignment (MSA) was used to generate a Hidden Markov Model (Eddy 1996). The sequences generated were subsequently used to generate a new alignment in the MUSCLE server with default conditions (Figure 6A, step *a*), or to collect remote homologous sequences from the UNIPROT database using PSI-BLAST that were aligned using MUSCLE (step *b*). The final alignment including proteins from all kingdoms was exclusive to Na^+^/H^+^ exchangers related to AtNHX1, and highly reliable to infer position-specific evolutionary information for this transporter. A final step was done using only the AtNHX1 PDB sequence generated by SwissModel and the ConsurfServer (http://consurf.tau.ac.il/) default conditions to generate the MSA for the conservation analysis (Figure 6A, step *c*). Based on the MSA obtained in steps *a* and *b* (formed by 352 and 243 sequences respectively), or the MSA defined by the ConsurfServer-sequence, evolutionary conservation scores were calculated using a Bayesian method (Mayrose et al. 2005), and the ConSurf web-server (http://consurf.tau.ac.il/) (Landau et al. 2005). The scores obtained were projected onto the 3D model of AtNHX1.

**Figure 6.**
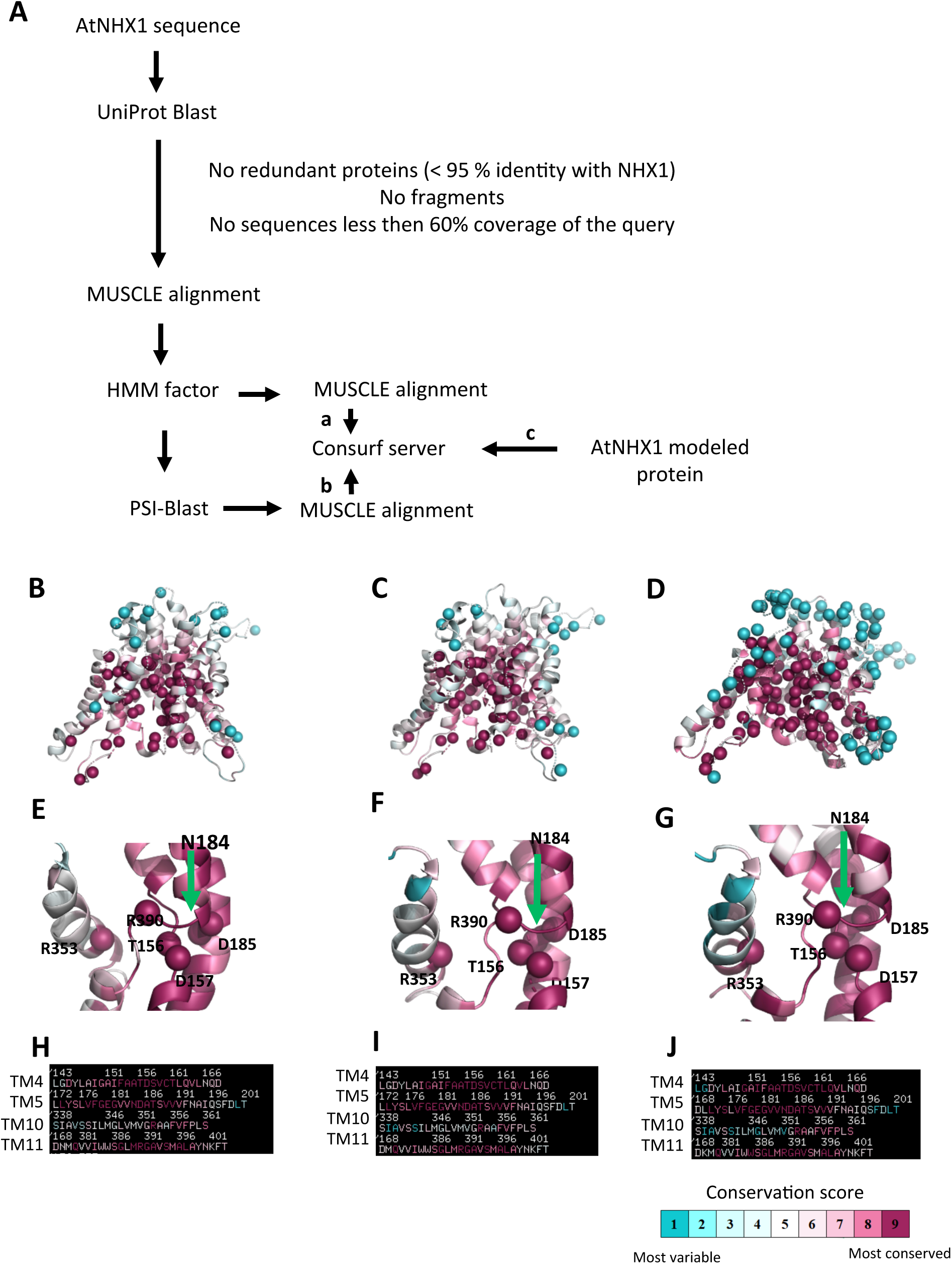
Evolutionary conservation of AtNHX1. **(A)** Flow chart of the methods followed to obtain the evolutionary conservation profile of AtNHX1 calculated via the ConSurf web-server (http://consurf.tau.ac.il). **(B-G)**. The evolutionary conservation profile of AtNHX1 obtained by the previous methods; **(B)** and **(E)** corresponds to the process following arrow *a*; **(C)** and **(F)** with arrow *b*, and **(D)** and **(G)** with arrow *c*. The profiles are colored according to their conservation-grades using the color-coding bar, indicating turquoise the most variable residues and purple the most conserved. The most variable and most conserved positions of each transporter are shown as spheres. In all three cases the pattern was repeated: a highly conserved intramembranous core while intervening loops and lipid-facing residues are variable. **(H-J)** Representation of the sequences in TM4, 5, 10 and 11 for each process following arrows routes *a*, *b* and *c* respectively. Note the high conservation of the residues in TMs contributing to the Nha-fold in the active center. TM10 is not highly conserved, except for the R353 residue. The AtNHX1 model shown is based on PaNhaP template.

The results showed that the model structure of AtNHX1 is compatible with the conservation pattern: the protein hydrophobic core is highly conserved while the residues facing the lipid bilayer or located in extramembranous regions are variable (Figure 6B, C and D). These results are similar to the previously described for PaNHX3, HsNHE2, and HsNHE9 (Kondapalli et al. 2013; Schushan et al. 2010; Wang et al. 2014). Of note is the high conservation observed among the residues found in the core of protein near the active center, and more importantly in TM4, TM5 and TM11 (Figure 6H, I, J). These sites are highly occupied by titratable residues, whose presence in the membrane is often associated with function. It is also noteworthy that although the overall conservation of TM10 is mostly low, residue R353 shows a high conservation score (Figure 6E, F and G). Interestingly, variability of peripheral amino acids seems to be higher at the vacuolar side of the membrane than the cytosolic side.

### Conserved residues with functional roles

The results described so far have shown the high structural conservation among the CPA superfamily in general, and more specifically among the CPA1 family members, making it plausible to translate the knowledge obtained from prokaryotic proteins to better understand the function and structure of eukaryotic proteins. However, it is important to note the need to experimentally confirm the conclusions since even between phylogenetically close proteins structural differences can be observed that in turn could result in different mechanism of transport or protein regulation (Masrati et al. 2018). To determine the relevance of the conserved amino acids described above in AtNHX1 activity, site-directed mutations were done at each residue judged as structurally important and the resulting protein variants were tested for activity in the yeast *Saccharomyces cerevisiae* as reported previously (Hernandez et al. 2009; Quintero et al. 2000).

The yeast strain AXT3K lacks the plasma membrane Na^+^ efflux proteins ENA1-4 and NHA1, and the prevacuolar Na^+^/H^+^ exchanger ScNHX1, which renders these cells sensitive to NaCl and hygromycin B (HygB) (Quintero et al. 2002). The expression in AXT3K cells of the AtNHX1 and AtNHX2 proteins suppressed the salt- and hygromycin-sensitive phenotype of this strain (Barragán et al. 2012; Quintero et al. 2002; Yokoi et al. 2002). Hence, the wild-type AtNHX1 and the mutant alleles were cloned in the yeast expression vector pDR195 and transformed in the yeast strain AXT3K to test the functionality of the mutant proteins in both solid and liquid media. Plasmid pDR195 uses the strong and mostly constitutive *PMA1* gene promoter to drive the expression of the recombinant protein. Expression of wild-type AtNHX1 improved AXT3K growth in the presence of 50 µg/ml of HygB (Figure 7). Cells transformed with the mutant alleles showed different phenotypes. The mutation of the D185 residue into a Leu or Asn generated non-functional proteins, as expected since D185 belongs to the active center residue of CPA1 proteins. Other mutations of residues at the core of the active center of the protein that are part of the Nha-fold (D157N, R353L) generated equally inactive proteins (Figure 7). These results indicate that the conserved residues of the active core in AtNHX1 have an important functional and/or structural role in the protein, and they are essential for protein activity in the conditions tested. However, the mutation in which Asp substituted for N184 (N184D) in the active center suppressed the sensitivity to HygB partly (Figure 7). This result implies that the N184 is important but not essential for activity.

**Figure 7.**
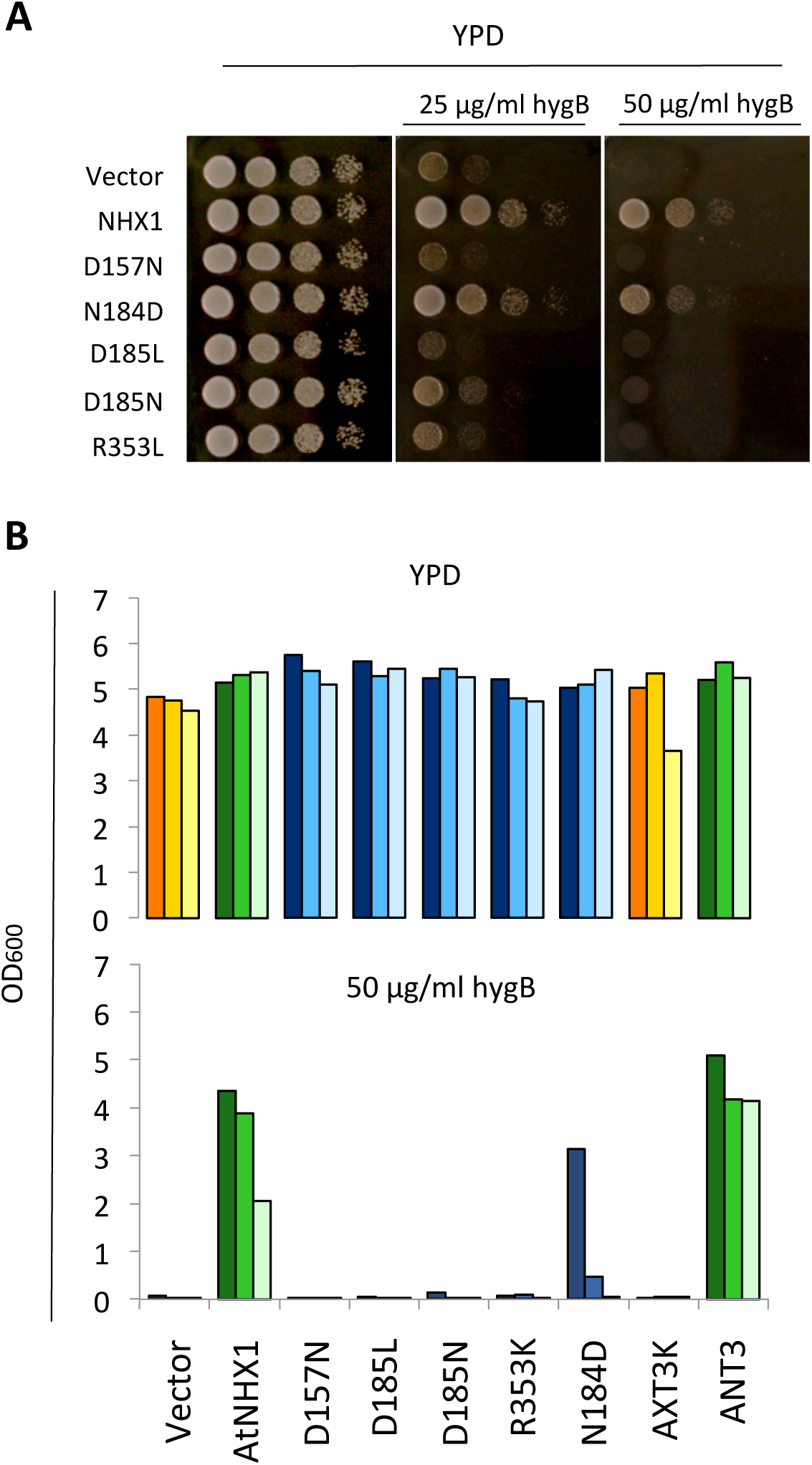
Complementation of the yeast AXT3K strain with an allelic series of AtNHX1 mutants in conserved residues. The cDNAs of AtNHX1 and of the indicated mutant alleles of conserved residues were cloned into the yeast expression vector pDR195 and transformed into strain AXT3K (*Δena1-4 Δnha1 Δnhx1*). Overnight cultures were normalized in water to OD600 of 0.5. Aliquots (5 μL) from normalized cultures and 10-fold serial dilutions were spotted onto YPD solid medium plates **(A)** or used to inoculate 200 μL of YPD liquid medium in 96-well plates **(B)** with different concentrations of hygromycin B. Liquid cultures were grown at 30°C overnight before measuring the optical density. Pictures were taken after 2-3 days at 30°C.

Previous reports have shown that the antiporters HsNHA2 and EcNhaA lost their transport activities when the DD-motif of CPA2 proteins was substituted for an ND-motif (Lee et al. 2013b; Schushan et al. 2010). However, the mutation N184D to generate a DD-motif only decreased AtNHX1 protein activity, but it was still able to overcome the HygB sensitivity (Figure 7). Thus, the integrity of the ND-motif is not essential for AtNHX1 activity, but it is important to reach optimal functioning. Recently Uzdavinyz et al (2017) have demonstrated that the electrogenic properties of CPA2 family members is not due to this DD-motif as previously thought, but to the presence of a conserved Lys in TM10, which in CPA1 members is exchanged for an Arg that promotes electroneutral activity (Figure 1B). This position in AtNHX1 is occupied by R353, which is coherent with the electroneutral exchange of AtNHX1 (Gradogna et al. 2021; Venema et al. 2002). Mutagenic studies of this conserved Arg in TM10 in different CPA1 family members have been performed but results were inconclusive (Hellmer et al. 2003; Schushan et al. 2010; Wang et al. 2014). To study the relevance of the conserved R353 in TM10 of AtNHX1, and to explore whether it was possible to make AtNHX1 electrogenic, a set of allelic variants was generated at the R353 position, so that all possible conditions of ND/DD motifs and R/K residues were studied: DD-Motif + R353/R390; ND-motif + R353K; ND-motif + R390K; DD-motif + R353K; and DD-motif + R390K. In this allelic series the R390K mutation was used as control to determine whether the electrogenicity (if changed) was dependent only of residue R353 or the active center motif ND/DD.

Functional assays using the AXT3K yeast strain would not directly allow determining whether AtNHX1 was transformed into an electrogenic protein, but at least it would inform whether the new arrangement of catalytic amino acids was compatible with protein activity in vivo. As shown in Figure 8A, any allele bearing an R-to-K mutation generated an inactive protein in solid media conditions, irrespective of the presence of an ND- or DD-motif in the active center. The only mutant that remained active was the N184D single mutant, although with slightly less activity than the wild-type AtNHX1 protein as monitored by yeast growth. These results meant that N184 is not essential for the activity of AtNHX1, but the presence of conserved arginines R353 in TM10 and R390 in TM11 were indispensable to maintain protein functionality. Moreover, although a charged amino acid in the active center in TM5 is necessary for the activity, the positively charged N184 is not essential and it can be exchanged for a negatively charged amino acid like an Asp (N184D mutation) with minimal effects.

**Figure 8.**
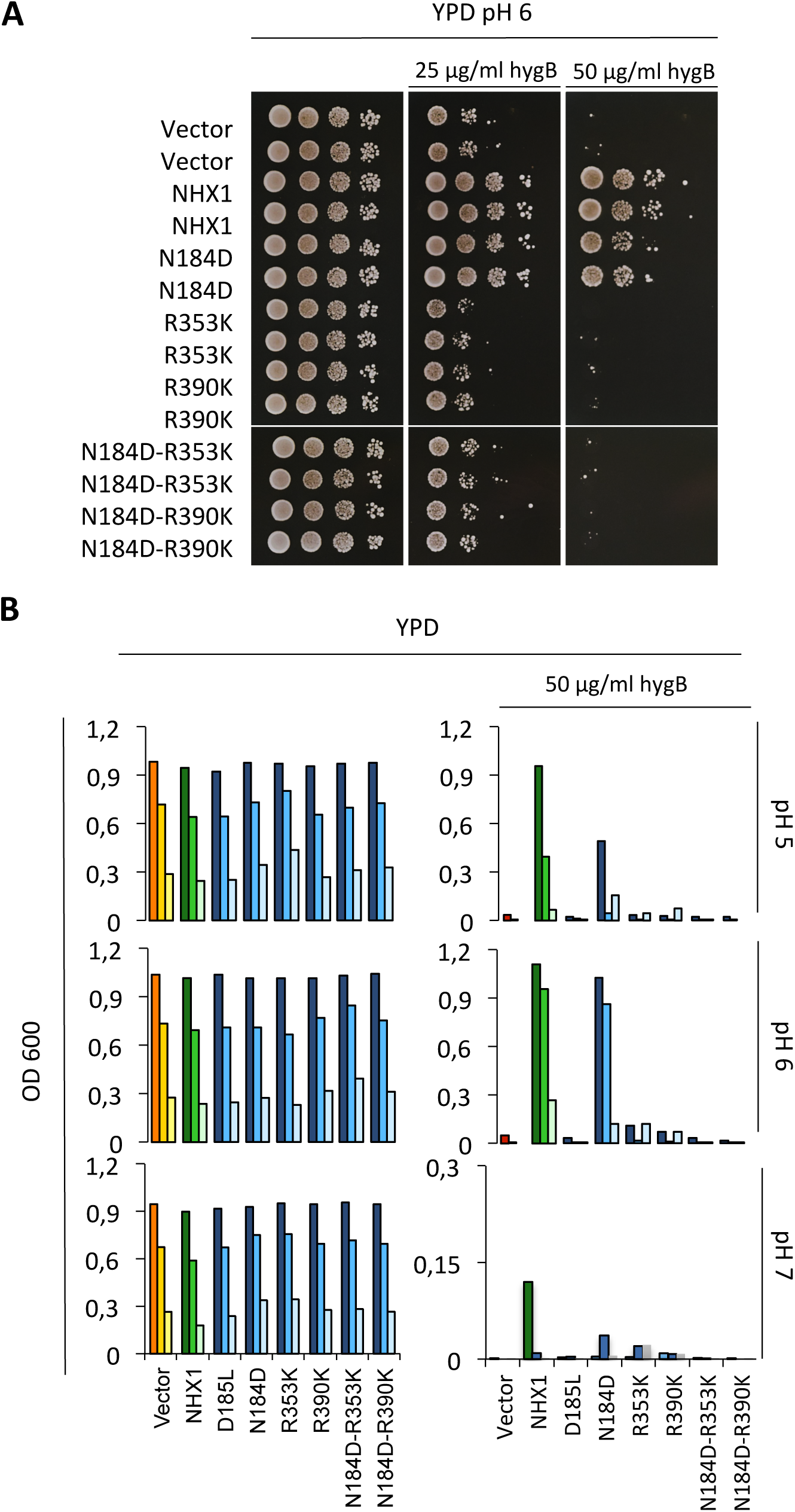
Functional assay of AtNHX1 mutant alleles in amino acids at the ion coordination pocket. The cDNAs of wild-type AtNHX1 and the indicated mutant alleles of the conserved residues were subcloned into the yeast expression vector pDR195 and transformed into the AXT3K (*Δena1-4 Δnha1 Δnhx1*). Overnight cultures were normalized in water to OD_600_ of 0.5. Aliquots (5μL) from normalized cultures and 10-fold serial dilutions were spotted onto YPD solid medium plates **(A)** or used to inoculate 200 μL of YPD liquid medium in 96-well plates with different concentrations of hygromycin B **(B)**. To determine whether the mutated amino acids had a role in pH sensing, the experiment in liquid media was performed at different pH (buffering with 10 mM MES). The liquid cultures were grown at 30°C overnight before measuring the OD. Plates were incubated 2-3 days at 30°C and pictured.

In the CPA2 protein family there is an interaction between a conserved Lys in TM10 and the Asp residue of TM5 in the active center. This Lys does not participate in ion binding; rather a competition-based transport mechanism has been suggested. The exchange cycle would start with a periplasmic/outer side open conformation of the protein in which D164 (in EcNhaA) or D157 (in TtNapA) is unprotonated, whereas D163 and D156, respectively, are engaged in a salt bridge with K300 or K305, which stabilizes this conformation. At low H^+^ concentration, the binding of ions to the DD-motif breaks the salt bridge, the structure becomes less rigid, and a conformational transition takes place releasing the transported ions to the cytosol. Deprotonated D163 and D156 again form a salt bridge with Lys at TM10, inhibiting the reorientation of the unloaded transporter. When Na^+^ binds from the cytoplasm, the salt bridge is broken and a conformational transition allows Na^+^ release at the periplasmic/outer side of the membrane (Călinescu et al. 2017; Lee et al. 2014; Maes et al. 2012). The same mechanism has been described for PaNhaP and MjNhaaP1 (Călinescu et al. 2016). However, in MjNhaP1 and PaNhaP, the Arg replacing this Lys does not interact with the ND-motif but forms an ion bridge to the neighboring conserved glutamate in TM5 (Paulino et al. 2014) while the conserved Asn in TM5 interacts with the conserved Thr in TM4. The main feature of this competition-based transport mechanism is that it is self-regulatory, ensuring that transport activity is switched off at extreme pH values to prevent excessive acidification or alkalinization of the cytoplasm (Călinescu et al. 2016). This is in accordance to the notion that the K300 in EcNhaA and K305 in TtNapA seem to have more than a functional role in the protein, and that they could be part of the pH activation mechanism, as the pH dependence of both in K300R and K305R mutants were shifted to the alkaline side by one pH unit in comparison to the wild-type proteins (Călinescu et al. 2017; Kozachkov et al. 2007; Maes et al. 2012).

Based in these considerations, we aimed to determine whether the non-functional mutants of AtNHX1 in medium YPD plus HygB had a different behavior under other growth conditions. To that end, similar assays were performed in liquid YPD media buffered at different pH, with and without addition of HygB. Of note is that the inhibitory effect of HygB was much higher in the medium buffered at pH 7. Regarding the growth of yeast cells expressing mutant proteins, the results were similar in all tests: any of the alleles that included a R-to-K mutation (either in R353K or R390K) failed to recover the HygB-sensitive phenotype of AXT3K (Figure 8B) regardless of the external pH. There were also no differences in their growth without the antibiotic. Interestingly, the only active allele was again N184D, but it did behave differently in the presence of HygB at different pHs. The N184D mutation showed to be detrimental for the protein activity at pH 5 and pH 7, as indicated by the reduced ability to suppress the sensitivity to HygB (Figure 8B). No difference between wild-type and N184D proteins in supporting yeast growth with HygB was found at pH 6. Growth was also tested in medium supplemented with Li^+^, a toxic analog of Na^+^. The N184D allele was able to sustain grow like wild-type AtNHX1, while the R-to-K mutants could not suppress the ion sensitivity (Supplemental Figure S4).

Combined, these results of yeast growth in various conditions suggest that N184 in the active center might contribute to pH sensing. This would be in accordance to what has been previously described for EcNhaA and TtNapA (electrogenic proteins with a DD-motif) that are more active at basic pH in comparison to MjNhaP1 and PaNhaP (with the ND-motif), which have been shown to be more active at acidic pHs (Călinescu et al. 2014; Uzdavinys et al. 2017).

### Vacuolar pH and AtNHX1 mutants

The activity of AtNHX1 and AtNHX2 *in planta* constitutes a leak pathway for vacuolar protons and together they contribute to control the vacuolar pH (Andres et al. 2014). The results presented herein made us question whether the mutation of conserved residues could have an effect on the activity of AtNHX1 and the regulation of the pH in the vacuolar lumen (pHv). To obtain a better insight on this possibility, the vacuolar pH (pHv) of the AXT3K yeast strain expressing the mutant alleles of AtNHX1 was measured. To assess the roles of AtNHX1 and the different variants in the pH regulation of the yeast, the ratiometric fluorescein-based pH sensitive dye, 2ʹ,7ʹ-bis-(2-carboxyethyl)-5-(and-6)-carboxyfluorescein (BCECF) was used. Ratiometric dyes have the distinct advantage of not being significantly affected by dye loading, cell size, or tissue morphology, unlike other non-ratiometric dyes. The excitation spectrum of BCECF is sensitive to pH, so it can be used as ratiometric pH indicator. This dye has an isosbestic point so that when it is excited with a 450 nm wavelength, the emission remains constant independently of the media pH. However, after excitation with a 490 nm wavelength the ensuing fluorescence emission increases proportionally to the pH of the solution. A single emission wavelength (535 nm) is measured at the two excitation wavelengths, and the ratio of the intensity emitted (I_490_/I_450_) provides a measure of pH. However, BCECF has a p*Ka* value near 7.1, and for that reason the I_490_/I_450_ ratio does not accurately estimate low pH values. Different approaches have been developed to overcome this problem (Brett et al. 2005b; Diakov et al. 2013). We applied the method proposed by James-Kracke (1992), which allows converting intensities of BCECF fluorescence to pH values between 4 and 9 by using the formula used to calculate intracellular calcium concentration from the fluorescence of Fura2, inverted after using with a set of buffers titrated at different pHs and in the presence of a protonophore (FCCP or CCCP) to generate the calibration curve. The inversion of the equation is needed because H^+^ binding to BCECF causes a decrease in fluorescence, whereas Ca^2+^ binding to Fura2 causes an increase in fluorescence. This type of calibration procedure allows BCECF to be used over a wider range of pH. BCECF localizes into the vacuole of yeast when introduced in its acetoxymethyl ester form (BCECF-AM). In previous assays, BCECF-AM has been used to measure the pHv of the *S. cerevisiae nhx1* mutant, to monitor responses to changing extracellular condition, to study how V-ATPase mutations affect its activity, and to examine the communication between organelles and extracellular media (Diakov and Kane 2010; Diakov et al. 2013; Martínez-Muñoz and Kane 2008; Tarsio et al. 2011).

Previous studies have reported pHv values between 5.5 – 5.9 in wild type *S. cerevisiae* strains (Coonrod et al. 2013; Diakov and Kane 2010; Diakov et al. 2013; Martínez-Muñoz and Kane 2008; Tarsio et al. 2011). Experiments using BCECF in yeast have demonstrated that overexpression of the ScNHX1 protein resulted in the alkalinization of the vacuole and, conversely, the lack of expression in the *nhx1* mutant produced acidification (Ali et al. 2004; Brett et al. 2005b). It is important to note that *in vivo* pH measurements are sensitive to the growth and metabolic conditions of the yeast cells themselves, which can be a source of variability between measurements (Diakov and Kane 2010; Diakov et al. 2013; Tarsio et al. 2011). Moreover, it has been reported that pHv of wild-type and mutants rapidly change according to the external pH (Brett et al. 2011; Plant et al. 1999). Under acidic pH stress pHv in wild-type was around 5.28±0.14, and upon alkali stress the pHv jumped to 5.83±0.13. To avoid that experimental noise, pHv measurements in our experiments were taken from yeast cells at the same growth phase (early logarithmic phase) and all media were buffered to 6.0 with arginine. Fluorescence ratios were transformed in pHv values following the protocol described by James-Kracke (1992).

Firstly, yeast strains differing in the presence or absence of NHX exchangers were compared. Mutation of the *nhx1* gene produced vacuolar acidification of 0,83 pH units relative to the wild-type strain and expression of AtNHX1 restored wild-type values (Figure 9A). These results validated the experimental approach. To test whether the AtNHX1 mutants affecting conserved residues at the pore domain had an effect in the pHv, the BCECF experiment was repeated three times, using three independent colonies each time. The results comparing the pHv of yeast cells expressing various AtNHX1 mutants showed a consistent tendency in independent experiments, but experimental variation in absolute pHv values between experiments meant that no statistically significant differences could be demonstrated in most samples. The lack of statistical significance could be due to the dynamic range of the ratiometric dye or the calibration method. Representative data from one of the experiments is shown in Figure 9B. As expected the dead-protein mutant D185L had an acid pHv similar to the empty-vector control since the protein is inactive. On the contrary, the N184D mutant had a near wild-type pHv, demonstrating again that the N184D mutant protein behaves as wild-type in medium buffered at pH 6.

**Figure 9.**
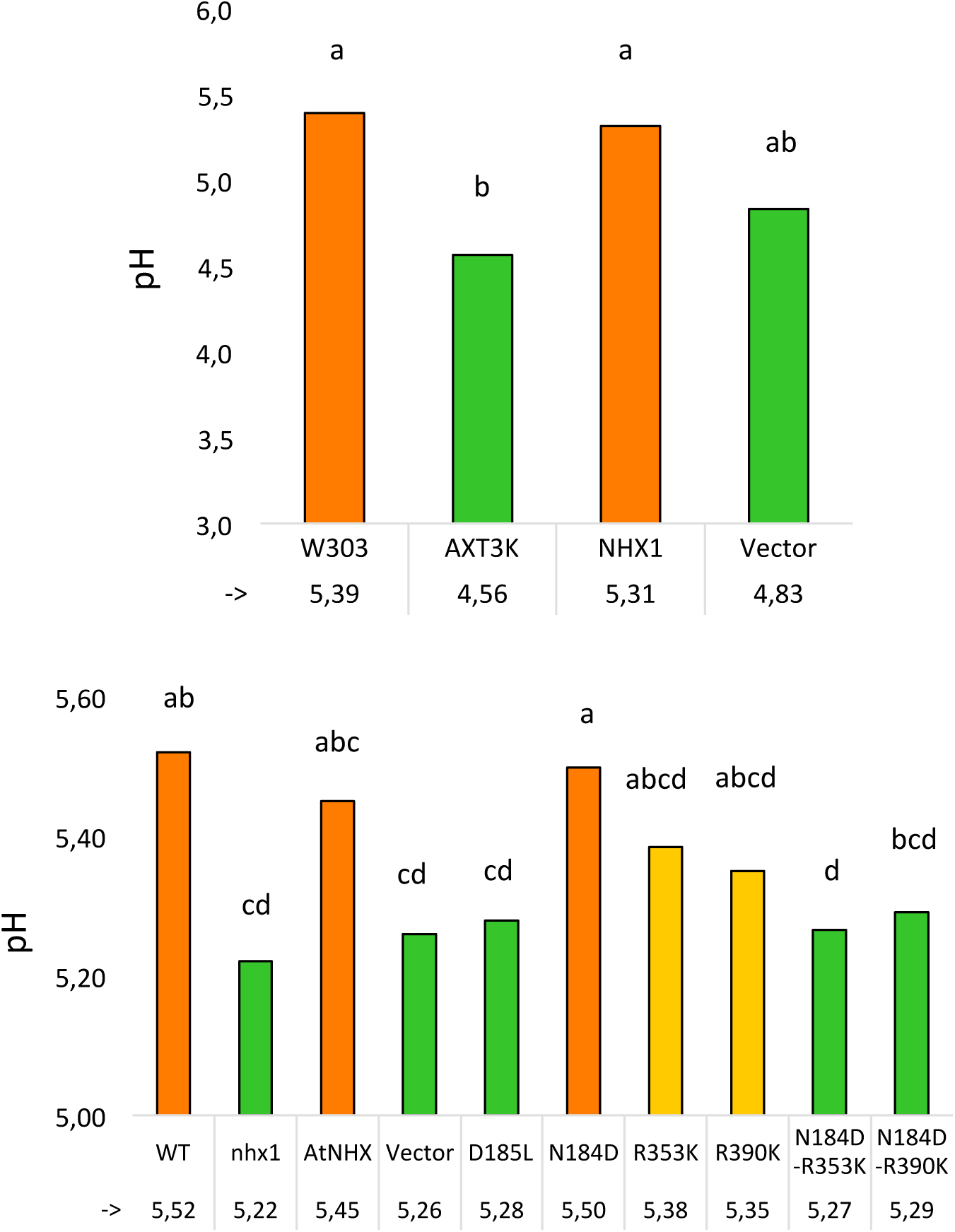
Vacuolar pH measurement in yeast expressing the AtNHX1 mutants by BCECF-AM fluorescence. **(A)** Vacuolar pH of the wild type (W303) and mutant (AXT3K) strains. AXT3K vacuolar pH was measured in the untransformed strain, and transformed strains with the empty vector or expressing AtNHX1. **(B)** Vacuolar pH values of the AtNHX1 mutant alleles. In both graphics the results of one representative experiment is represented. In each experiment three independent colonies were used to measure the vacuolar pH. Different letters indicate statistically significant differences in pairwise comparison by the Tukey HSD test (p< 0.05)

Surprisingly, conservative mutations in the conserved arginines R353K and R390K showed intermediate pHv values halfway between the wild-type and the null mutant D185L and empty vector controls, suggesting compromised exchange activity and reduced H^+^ leak. This result is somehow in contrast with the null-mutant phenotype showed by these mutants in the HygB and Li^+^ tolerance test (Figure 8 and Supplemental Figure S4). However, combining the R353K and R390K mutation with the N184D mutation that had no effect per se, produced acidic vacuoles similar to loss-of-function mutants.

## DISCUSSION

### Generation of topological and tridimensional models based on phylogenetic relatedness

The NHX exchangers of plants are secondary transporters of the cation-proton antiporter (CPA) family (Brett et al. 2005a; Rodriguez-Rosales et al. 2009). CPA antiporters are conserved across all biological kingdoms and have essential roles in pH, ion homeostasis and volume control. CPA1 antiporters are electroneutral and exchange one cation (K^+^ /Na^+^) against one H^+^ (Călinescu et al. 2014; Wöhlert et al. 2014). Among these proteins, in which plant NHXs are included, are also found proteins PaNhaP, MjNhaP1 and HsNHE1. CPA2 antiporters, including EcNhaA from *E. coli* and TtNapA from *Thermus thermophilus*, are electrogenic, exchanging one Na^+^ against two H^+^ (Hunte et al. 2005; Lee et al. 2013b; Uzdavinys et al. 2017). This family also contains eukaryotic proteins, e.g. HsNHA2 and the plant CHX clade (Brett et al. 2005a; Schushan et al. 2010; Sze and Chanroj 2018).

The first atomic structure of a cation-proton antiporter (CPA family) was the Na^+^/H^+^ exchanger of *E. coli* NhaA (EcNhaA). The 6 Å structural map of EcNhaA resolved 12 transmembrane helices (TMs) with a characteristic folding of the active center, named the Nha-fold, consisting in two unwound transmembrane stretches that cross each other in the middle of the membrane near the ion binding site by their unwound region creating an X-shaped structure. Later, the structures of two CPA1 members, MjNhaP1 and PaNhaP, were also obtained. Although they have some functional and structural differences with EcNhaA, the overall structure is well conserved, including the Nha-fold of the active center. The information gained from these structures allowed the *in silico* modeling and a better understanding of the architecture of the more complex eukaryotic CPA proteins, such as eukaryotic human HsNHE1 and HsNHA2 (Lee et al. 2013a; Schushan et al. 2010), *Populus euphratica* PeNHX3 (Wang et al. 2014), and of *A. thaliana* AtCHX17 (Czerny et al. 2016) and AtNHX6 (Ashnest et al. 2015). Most modeled structures were based in the first structure available, that of EcNhaA, even if it was not the most adequate model since these CPA1 proteins have been modeled using as template a CPA2 protein (Lee et al. 2013a; Wang et al. 2014). In our work with AtNHX1 four structures available at the SwissModel database were selected as templates because they fitted best to the AtNHX1 protein sequence. These four templates were *Methanococcus jannichii* NhaP1 (MjNhaP1) (Paulino et al. 2014), *Pyrococcus abysii* NhaP (PaNhaP) (Wöhlert et al. 2014), *Thermus thermophilus* NapA (TtNapA) (Lee et al. 2013b), and *E. coli* NhaA (Padan et al. 2009). We used these proteins as the reference structures to generate our own model of a NHX protein that was also compared to previous modeling reports. The alignment of the primary sequence of AtNHX1 with each of these four proteins individually and the alignment of all proteins together made evident the high conservation of the core structure of the family. Even though EcNhaA showed the lowest sequence conservation at the N- and C-termini among the template proteins compared, the main core in the pore domain of the proteins remained highly conserved, as it had been noticed before in the modeling of HsNHE1 and PeNHX3 proteins (Schushan et al. 2010; Wang et al. 2014).

The 3D structure obtained for AtNHX1 with each of the model templates were highly similar, with the main differences being in a more lax or tight structure of the protein, which in turn was reflected in the possible interactions that were predicted to take place among residues in the active site of the protein. However, the general structure was the same in all cases, and the predicted transmembrane segments based in individual alignments or in the overall alignment overlapped in most cases. This allowed to generate a topological model that was similar to the previously described for the eukaryotic HsNHE1 or PeNHX3, with 12 TM segments in the hydrophobic N-terminal part, or pore domain, and a long hydrophilic C-terminal tail, with both protein ends being cytosolic. Previous models to explain AtNHX1 topology were inconsistent. Yamaguchi et al. (2003) proposed a model according to which AtNHX1 topology comprised nine transmembrane domains, three hydrophobic regions not spanning the tonoplast membrane, and with the hydrophilic C-terminal domain in the vacuolar lumen. However, this topology is incoherent with a protease protection assay showing that the C-terminal tail of AtNHX1 was cytosolic (Hamaji et al. 2009), and with he proposal that AtNHX1 and HsNHE1 share the same topology, except that the first N-terminal TM segment of HsNHE1 is a signal peptide missing in AtNHX1 (Sato and Sakaguchi 2005). However, the major disadvantage of the approach followed by Sato and Sakaguchi (2005) was that the method used to obtain the topological model dealt with fragments of the protein instead of the entire protein. Based in a sequence and structure conservation approach, the model proposed herein yields a topology that is consistent with the common topology model proposed for the CPA superfamily: 12 TM antiparallel segments with a hydrophilic and cytosolic C-terminal tail. An important corollary of this topology is that the interaction of AtNHX1 (and AtNHX2) with CML18 cannot take place in the vacuolar lumen as reported by Yamaguchi et al. (2003).

While this work was in progress, the modeled structure of AtNHX1 generated using the AlphaFold AI software was released (Jumper et al. 2021). The predicted structure was aligned with the models previously created using the Swiss-model repository (Supplemental Figure S5). The alignment between the models confirmed the presence of the Nha-fold in the active site of the AtNHX1 protein, with the side chains of the conserved amino acids directed toward the intramembranous pocket in the center of the protein. The hydrophobic pore domain of the protein consists of TM segments arranged in a similar antiparallel fashion. However, the 3D structure generated by the AlphaFold software shows that the C- and N-terminal ends of the protein are located in different compartments of the cell. This is a direct consequence of the two complete TM segments (TM1 and TM2) modeled by the AlphaFold software, which in our model these segments are considered as two intramembranous (IM) semi-helices. However, these differences should not affect protein function or regulation. The final orientations of the other TM segments do not change compared to the topology predicted in this manuscript, and there are no previous reports of regulatory or activity domains in this region. All in all, the predicted 3D-structures aligned with a high level of accuracy, especially for those models generated with other CAP1 proteins templates.

### Validation of structural models

The best-scored tridimensional models of AtNHX1 using the SwissModel structural database and modeling tools were MjNhaP1 and PaNhaP. This is not surprising since the common ancestors of the CPA1 family are conserved membrane NhaP-like proteins from thermophilic bacteria and archaea (Brett et al. 2005a; Chanroj et al. 2012). Quality checks of the modeled structures based in different conventional analysis are essential to validate the model generated *in silico*. These analyses include the distribution of positively charged and hydrophobic residues (Fleishman and Ben-Tal 2006) and the pattern of evolutionary conservation (Bowie 2005). In the AtNHX1 models generated by MjNhaP1, PaNhaP and TtNapA, as most of the integral membrane proteins, AtNHX1 presents a higher number of positively charged residues (Arg and Lys) in intracellular positions compared with extracellular regions (Wallin and von Heijne 1998), while the polar residues are preferentially located in the intra- membrane core structures or in extra-membrane loops, and away from transmembranes at the periphery of the protein. Finally, the core residues are more conserved than those facing the lipids or those that are cytosolic or luminal (Fleishman and Ben-Tal 2006). These analyses have also been used to validate the predicted structure in other CPA modeling approaches (Ashnest et al. 2015; Lee et al. 2013a; Schushan et al. 2010; Wang et al. 2014). The AtNHX1 models generated using the MjNhaP1 and PaNhaP templates fitted perfectly the expected outcomes of these validation assays, and even the prokaryotic TtNapA, which belongs to the CPA2 family and has been used to model other CPA2 structures, e.g. AtCHX17 (Czerny et al. 2016) produced satisfactory results. However, the EcNhaA not only did not fit with the quality checks, but also one transmembrane segment was missing in the generated 3D structure (Supplemental Tables S11 and S2).

### Identification of conserved residues with functional roles

Integrating the primary sequence comparisons and the topological and ternary models, several residues with probable structural and functional significance were detected. The graphical representation of the TM segments in which these residues are located made evident that AtNHX1 present the Nha-fold at the active center as previously demonstrated for crystalized proteins of the CPA superfamily (Lee et al. 2013b; Padan et al. 2009; Paulino et al. 2014; Wöhlert et al. 2014). The active center N184-D185 motif of the CPA1 electroneutral proteins is located in TM5 of AtNHX1. In the vicinity of the ND-motif the expanded sectors of TM4 and TM11 cross each other, and in these sectors the conserved T156-D157 (TD-motif) of TM4, and R390 of TM11 are located. Moreover, TM10 is in the surroundings of the Nha-fold and the conserved R353 in this TM can be found interacting with conserved residues in TM4 and TM11 (Figure 3).

The predominant view has been that the presence of the DD- or ND-motif in the active center of the protein determined the electrogenicity of the antiporter (Hunte et al. 2005). The additional negatively charged Asp residue in the DD-motif of CPA2 proteins was proposed to be responsible for the extra H^+^ that is transported by these exchangers compared with CPA1 members carrying the ND-motif (Călinescu et al. 2014; Hunte et al. 2005). For both EcNhaA and TtNapA the DD-motif is essential for the protein activity, and introducing an Asn residue in this position rendered the protein inactive (Inoue et al. 1995; Lee et al. 2013b). In CPA1 protein members, the substitution of the Asp residue in the ND-motif showed that it is essential for the protein activity. However, the Asn residue was not. Replacing N160 in MjNhaP1 for an Ala generates an inactive protein, but the N160D substitution mutant of this protein (Hellmer et al. 2003), as well as the N187D substitution in PeNHX3 generated active proteins (Wang et al. 2014). Similar results were obtained for AtNHX1 since the N184D mutation also produced a biologically active protein, whereas mutations D185L and D185N produced inactive proteins (Figure 7). Nonetheless, Asn to Asp mutations in the active enter of CPA1 proteins do not render these proteins electrogenic (Hellmer et al. 2003; Paulino et al. 2014; Wang et al. 2014). In the case of AtNHX1 more analyses are required to confirm this. The N184D mutant showed to be detrimental for the yeast growth in the presence of hygromycin, which is indicative of compromised activity, but supported a slightly more robust growth than the wild-type AtNHX1 when grown in 10 mM LiCl (Figure 8 and Supplemental Figure S4). These results indicate that the highly conserved N184 in the active center is not essential for the activity of AtNHX1, as previously demonstrated for MjNhaP1 or PeNHX3 (Paulino et al. 2014; Wang et al. 2014), but could have a role in substrate selectivity by changing the geometry of the active center. In MjNhaP1, it was proposed that the homologous Asn is necessary to stabilize the proton or substrate-bound state, which can also be fulfilled by Asp (Paulino et al. 2014). As for HsNHA2, the mutation of de DD-motif into an ED-motif decreased Li^+^ tolerance, which is opposite to what we observed when the ND-motif of AtNHX1 is exchanged for a DD-motif (Supplemental Figure S4), implying again that the presence of an Asp in the first position of the DD motif can modify ion affinity and/or selectivity (Schushan et al. 2010).

Residue R353 in AtNHX1 located in TM10 has counterpart homologues in MjNhaP1 (R320) and PaNhaP (R337), both in TM10. The X-ray structure of these proteins shed light on their function in these proteins. These Arg residues form ion bridges with a conserved Glu in TM5: R320 with E156 in MjNhaP1, and R337 with E154 in PaNhaP. This Arg-coordinating Glu is conserved in all CPA1 antiporters, and the corresponding residue in AtNHX1 is E180. These highly conserved residues essential for activity seem to have a role in stabilizing the protein (Goswami et al. 2011; Hellmer et al. 2003). Recent studies showed that the electrogenic activity of TtNapA is due to the presence of a Lys in TM10 (K305) but not to the DD-motif in the active center (Uzdavinys et al. 2017). This Lys is the equivalent to residue to R353 of AtNHX1. They also demonstrated that HsNHA2 is an electroneutral protein despite of the DD-motif in the active center and because of the presence of an Arg (R432) in the homologous position to K305 of TtNapA. Later, Călinescu et al. (2017) showed that, although having an important role in protein activity and stability, the homologue Lys in EcNhaA (K300) is not essential for electrogenic transport. Together, these results indicate that the DD-motif is not the basis for electrogenicity and that the K/R dichotomy in TM10 explains only partially this transport mode.

In the CPA2 family proteins there is an interaction between the conserved Lys in TM10 and the Asp residue in the active center, and this interaction regulates the pH sensing and the activity of the protein in a competition-based transport mechanism that ensures transport activity as long as no extreme pH values are reached, in order to prevent excessive acidification or alkalinization of the cytoplasm (Călinescu et al. 2017; Călinescu et al. 2016; Lee et al. 2014; Maes et al. 2012). In the case of PaNhaP and MjNhaP1, the Arg replacing the Lys does not interact with the ND-motif but forms an ion bridge to the neighboring conserved glutamate in TM6, and the conserved Asn of the ND-motif in TM5 interacts with a conserved Thr in TM4. The essential role of R353 in protein function or stability has been confirmed by functional analysis in yeasts (Figures 7 and 8). In the mammalian CPA1 proteins NHE1 and NHA2, the Arg in TM10 is conserved (Figure 1B) but, unlike the AtNHX1 mutant R353K, the R-to-K mutant of NHA2 retained partial activity. Something similar occurs with the conserved positively charged residue in TM11 (R458 in HsNHE1, K460 in HsNHA2, R390 in AtNHX1). The mutant K460A of HsNHA2 exhibited a Li^+^ selective phenotype signifying that the mutant protein had gained greater affinity for Li^+^ than the wild-type protein (Landau et al. 2007; Schushan et al. 2010). In prokaryotes, the functionally important Arg in the unwound TM11 (R285 in MjNhaP1, R362 in PaNhaP, R390 in AtNHX1) might have as well a stabilizing function by interacting with residues in the active site (Paulino et al. 2014; Wöhlert et al. 2014), but not much is known about them. Our data show that even the conservative change R390K was unable to recover the AXT3K sensitivity to HygB (Figure 8) indicating that R390 might have as well a functional or structural role in the AtNHX1 protein.

In EcNhaA the conserved residues T132 and D133 (TD-motif) are involved in the ion coordination and translocation, but they are not essential for protein activity (Galili et al. 2002; Maes et al. 2012). Instead, a function in the stabilization of the active site has been proposed (Maes et al. 2012) and recently confirmed (Rimon et al. 2018). In the archea proteins PaNhaP and MjNhaP1 the TD-motif is conserved and is essential for the coordination of the ion (Paulino et al. 2014; Wöhlert et al. 2014). In PaNhaP and MjNhaP1 residues Thr129 and Thr131 have been described to interact by their side chain with N158 and N160, respectively, which do not participate in the ion coordination but control the access to the ion-binding site. In the mammalian NHE1 and NHA2, and in TtNapA this TD-motif is not conserved. Although for PeNHX3 only the presence of a non-charged Tyr in position 149 was described (Wang et al. 2014), the alignment of the CPA proteins performed herein indicates that the TD-motif is in fact conserved (Figure 1B). This motif is also conserved in AtNHX1 (T156-D157), and according to all the models obtained for AtNHX1, there seems to be an interaction of T156 with N184, as described for the archea NhaP proteins. Our yeast complementation assays showed that D157 of the TD-motif is essential for AtNHX1 activity, but a mutant in T156 was not tested.

The AtNHX1 topological and tridimensional structure generated by homology modeling in this study sheds light on its transport mechanism. In addition, the analysis of the most conserved residues demonstrate that AtNHX1 maintain essential properties of the CPA1 family, but also carries unique features in ion binding and translocation. Additional ion transport assays in tonoplast vesicles should be done to confirm the functionality of the described amino acids in the electroneutral nature of AtNHX1 or in the affinity traits.

### Regulation of vacuolar pH by AtNHX1

One of the main functions of NHX proteins is to regulate the luminal pH of organelles in which they reside (Andres et al. 2014; Bassil et al. 2011; Reguera et al. 2015). This function is highly conserved in eukaryotic organisms, including yeasts in the which *Δnhx1* mutant has been shown to have a more acidic endosomal pH when subjected to acid stress (Ali et al. 2004; Brett et al. 2005b; Diakov et al. 2013; Plant et al. 1999). The yeast ScNHX1 protein resides in late endosomes/pre-vacuolar compartments, and ScNHX1 effects on organelle pH seem to be tied to intracellular trafficking. The *nhx1* mutant is also known as *vps44* (vacuolar protein-sorting 44) (Bowers et al. 2000). The *vps44/nhx1* mutant secreted 35% of a vacuolar carboxypeptidase Y (CPY) and missorted markers associated with PVC, Golgi, or vacuolar membrane. These results evidence that (Na^+^,K^+^)/H^+^ exchange activity is essential for endosomal function and protein sorting in eukaryotic cells, and that proper pH homeostasis in LE/PVC is critical for the sorting of proteins.

When extracellular pH approaches the cytosolic pH, nutrient and ion uptake can be disrupted because the pH gradient across the plasma membrane is lost. The *ENA1* gene, encoding a Na^+^-ATPase, is induced by alkaline conditions and encodes a pump capable of exporting toxic Na^+^ in the absence of a H^+^ gradient (Haro et al. 1991). This protein is mutated in AXT3K, which may explain why reduced yeast growth was observed at pH 7.0 (Figure 8). The expression of AtNHX1 conferred yeast tolerance to high K^+^ or Na^+^ at acidic external pH, but on alkaline medium AtNHX1 was ineffective (Sze and Chanroj 2018). Attempts to monitor AtNHX1 in yeast based on resistance to hygromycin instead to Na^+^ or Li^+^ were also unsuccessful because, for unknown reasons, to toxicity of hygromycin increased greatly at pH 7 (Figure 8).

Measurements of luminal vacuolar pH (pHvac) in yeast cells by the pH-sensitive fluorochrome BCECF showed results that vacuoles of the *nhx1* mutant were more acidic than that of the wild type (Figure 9), which is consistent with the idea that ScNHX1 catalyzes K^+^ uptake and H^+^ efflux from compartments acidified by the vacuolar H^+^-pumping ATPase (Brett et al. 2005b). However, although trends in pHvac differences between genotypes were maintained, quantitative estimations of pHvac varied notably from experiment to experiment, complicating the statistical analysis of data. Cytosolic and vacuolar pH can vary significantly under different growth conditions and during different growth phases, which could explain these variations (Diakov et al. 2013). Even though preventive measures were taken, the source of the experimental noise when estimating the pHvac between experiments could not be cancelled. Nonetheless, qualitative differences in pHvac between genotypes marked clear tendencies regarding the contribution of NHX activity to pHvac that were coherent with the complementation test of AXT3K cells.

Important differences were observed in the pHvac of strain AXT3K expressing various AtNHX1 mutants affected in conserved residues. The vacuole of yeast carrying the AtNHX1 mutant N184D had a more basic pHvac than AXT3K and similar to the wild-type AtNHX1, which correlates with the ability of this mutant to suppress the sensitivity to HygB. In CPA1 proteins, the conserved Asn in the active center (ND-motif) may interact with the TD-motif at the Nha-fold (Figure 3 and Supplemental Figure S3). The exchange of Asn by Asp in the N184D mutant of AtNHX1 might alter this interaction and maintain the protein active at the optimal external pH 6, but it seems to be detrimental for growth at pH 5 (Figure 8). The equivalent N187D mutant of PeNHX3 also complemented the yeast *nhx1* mutant, but the vacuolar pH was not measured (Wang et al. 2014). In MjNhaP1, the N160D mutant showed lower activity compared to the wild type (Paulino et al. 2014).

Surprisingly, single mutants with the two conserved arginines R353 and R390 changed to Lys showed a pHvac intermediate between the wild-type AtNHX1 and a loss-of-function mutant (Figure 9). This was an unexpected result since in the growth assays no complementation of the AXT3K sensitivity to HygB or LiCl by R353K and R390K mutants was detected. The double mutants in which the ND-R arrangement of AtNHX1 was completely changed to the DD-K configuration found in electrogenic CPA2 proteins showed an pHvac similar to loss-of-function mutants, e.g. D185L (Figure 9). The phenotype of yeast expressing the R-to-K variants could be explained by a lower activity of the AtNHX1 protein that was insufficient to suppress the phenotype of the yeast *nhx1* mutant but enough to constitute a H^+^ shunt preventing extreme acidification of the vacuolar lumen. Although in normal growth conditions these proteins are able to translocate K^+^ into the vacuole in exchange of H^+^, the selective conditions applied in the functional assays in yeast could still be inhibiting the yeast growth thereby masking the reduced protein activity. This low activity of R353K and R390K mutants could be explained by a change in ion selectivity or the inability to coordinate properly the substrates due to an altered, less than optimal structure. This could also explain the sensitivity of AXT3K cells expressing R353K and R390K mutant proteins in AP media with LiCl. Moreover, in the event that the nature of the mutant AtNHX1 protein changed from electroneutral to electrogenic, it could be deleterious to yeast in selective conditions even though the protein remained active. Nevertheless, all these suppositions are only based on indirect evidence gathered from growth assays and measurements of the vacuolar pH as affected by the AtNHX1 mutants. A direct assay measuring the ion transport of the mutants in tonoplast vesicles is necessary.

In TtNapA, the mutation of the conserved K305 for other basic amino acids, such as Gln or Arg, generated alleles only slightly active at pH 8. Moreover, the new alleles had lost electrogenic properties and their activity became electroneutral. The activity could only be recovered to wild-type levels in a D156N-K305Q mutant. This made evident the need of an interaction of a positive amino acid different from Lys in TM10 with the Asn in the active center for the activity of electroneutral proteins (CPA1). This double mutant was also electroneutral, and the mutation of K to R alone was enough for this conversion. Recently, it has been reported that EcNhaA could be converted from electrogenic to electroneutral by mutating the D163 in the DD-motif into Asn (ND- motif), together with two other amino acids different to the conserved amino acids herein mentioned, but with strategic positions near the active center (A106S and P108E) (Masrati et al. 2018). These results demonstrate that is not only essential the presence of certain conserved amino acids for the activity of the protein, but also the interactions they maintain to keep the conditions in the active site for the kind of transport that takes place. However, according to Uzdavinys et al (2017) the conserved Lys in TM10 is essential for the electrogenicity of the protein, while for Masrati et al (2018) it is the first Asp of the DD-motif. Nonetheless, in both cases it was possible to mutate the active site of an electrogenic DD-motif into a ND-motif obtaining an active protein, always balanced by mutations of other strategic residues that maintained the right conditions for the translocation of ions, generating electroneutral proteins from a electrogenic one. This would mean that to convert AtNHX1 into electrogenic would probably require more than one mutation. This has also been observed in previous attempts trying to create electrogenic proteins out of CPA1 members (Wang et al. 2014). Based in published results with other CPA proteins, the N184D-R353K mutant of AtNHX1 may act already as electrogenic, but transport assays in yeast vesicle should be done to demonstrate this point.

In conclusion, the topological and ternary models of AtNHX1 indicate that this plant protein conserves the structural features characteristic of microbial and mammalian members of the CPA superfamily. Several conserved amino acids that are essential for the activity of AtNHX1 have been identified at the active site and the Nha-fold. These residues are D157 of the TD-motif in TM4, D185 at the ND-motif in TM5, and arginines R353 and R390 in in TM10 and TM11, respectively. Residue N184 at the ND-motif that is highly conserved in electroneutral antiporters of the CPA1 family is not essential for the activity of AtNHX1, at least in the heterologous system used to validate the functionality of mutated AtNHX1 proteins. Last, in agreement with the proposed function *in planta*, AtNHX1 can regulate vacuolar pH in yeasts cells, and the expression of proteins mutated in the conserved amino acids of the Nha-fold generates pH differences in the yeast vacuole.

## MATERIALS AND METHODS

### Plasmid constructs

The mutant alleles of *AtNHX1* generated in this study are listed in Supplemental Table S3, together with the primers used to produce the point mutations by PCR amplification. To generate mutated alleles two different strategies were followed. In both cases the pBluescript (KS)-NHX1 plasmid was used as template. For the first set of mutants, the New England Biolabs’ Q5® Site-Directed Mutagenesis Kit was used, following the manufacturer’s specifications (https://www.neb.com, protocol E0554). Primers for this protocol were designed using the online tool supplied by NEB (http://nebasechanger.neb.com). The second set was obtained using a high-fidelity polymerase and the following protocol (Wang and Malcolm 1999). Primers for these reactions were designed such that they had the desired mutation flanked by 10 bp 5’ and 3’ fitting the template; and the reverse primer was the reverse complementary sequence of the forward primer. As both primer fit to each other better than to the template, two reactions were set, each one with only one of the primers, for 5 cycles. Afterwards, 10 μl of each reaction were mixed and, after addition of the polymerase, the combined reaction was allowed to proceed for 13 cycles. The elongation time was chosen to allow the whole template plasmid to be amplified. After the PCR reaction, 10 μl of the product was treated with 0.5 μl of *Dpn*I, which cuts only the methylated GATC sites, in the PCR buffer for 1 h 37°C. *E. coli* cells were transformed with 5 μl of the digestion. The generated alleles were subcloned in plasmid pDR195 as *Xho*I/*Not*I fragments and confirmed by sequencing. Yeast transformations were performed using the lithium acetate/PEG method (Elble 1992).

### Complementation and growth assays

Yeast growth assays were used to determine the tolerance of AXT3K yeast transformants to different NaCl or LiCl concentrations in AP medium, and different hygromycin B concentrations in YPD medium. Yeast were transformed with the pDR- NHX1 alleles and selected in YNB plates without histidine and uracil. Yeast transformants were inoculated in 2 mL of YNB media supplemented with the corresponding amino acids and grown at 30°C overnight. The cells were then harvested and resuspended to a final OD_600_ of 0,5. The cells were then serially diluted 10-fold in water, and 5 μl of each dilution were spotted onto the selective media: AP with different NaCl or LiCl concentrations, or YPD plates supplemented with different concentrations of hygromycin B. Plates were incubated at 30°C during 3-5 days.

For growth assays in liquid media, transformed yeast cells expressing the different pDR-NHX1 alleles were inoculated in 2 mL of YNB media supplemented with the corresponding amino acids and grown at 30°C overnight. The cells were then harvested and resuspended to a final OD_600_ of 0.5. Next, 20 μl of this dilution was used to inoculate a 96-well plate containing 200 μl of liquid media per well. From the first well, 10-fold serial dilutions were made. Plates were incubated at 30°C. The OD_600_ of the culture was measured 24 and 48 days after the inoculation with the Varioskan LUX Multimode Microplate Reader (ThermoFisher Scientific). Three independent AXT3K transformant colonies were used in each assay. The mean and standard error (SE) of the growth of each transformant was calculated using Microsoft Office Excel software.

### Measurements of vacuolar pH

To measure the vacuolar pH of yeast cells, the protocol described by Ali et al., 2004 was used, with modifications to make the pH measurements in a microplate reader (Brett et al. 2005b). Yeasts were grown overnight in AP pH 6.0 medium without histidine and uracil. Cultures in exponential phase of growth (OD_600_ 0.5-0.6) were harvested, washed two times and finally resuspended in AP medium without amino acids to a final OD_600_ of 0.2-0.3. Three independent colonies per mutant allele and controls were used. Each sample was incubated with 50 µm of 2’,7’-Bis-(2- carboxyethyl)-5-(6)-carboxyfluorescein acetoxymethyl ester (BCECF-AM; Molecular Probes, Eugene, OR) at 28°C, with gentle shaking. After 20 minutes, the culture was centrifuged 10’ at 5,000 rpm and washed three times with AP medium without amino acids and without BCECF-AM (incubating during 10 min with the new medium after each centrifugation). Finally, yeast cells were resuspended in 100 μl of AP medium without amino acids and without BCECF and the fluorescence was measured. Fluorescence intensity and absorbance values were recorded using a Varioskan LUX Multimode Microplate Reader (ThermoFisher Scientific) at 20 °C that were shacked before each measurement. Samples were sequentially excited by two wavelengths: 450 nm and 490 nm; emission fluorescence was detected at 535 nm for each of the two excitation wavelengths. Absorbance was also measured with 600 nm wavelengths. Three reads per well and wavelength were done. Measurements were repeated three times for each culture, washing the cells in between with AP medium without amino acids and without BCECF. At the end of each experiment, a calibration curve of fluorescence intensity versus pH was obtained. Samples were centrifuged and resuspended with 200 μL of the following calibration medium: 50 mM MES, 50 mM HEPES, 50 mM KCl, 50 mM NaCl, 200 mM ammonium acetate, 10 mM NaN_3_, 10 mM 2- deoxyglucose, 50 µM cyanide m-chlorophenylhydrazone. Buffers were titrated to eight different pH values (5.2, 5.6, 6.0, 6.4, 6.8, 7.2, 7.6, 8.0) using 1 M NaOH. Settings of the Microplate Reader were identical for measurements and pH calibration.

To estimate accurately acid pH values below 5.0, the approach of James-Kracke (1992) was followed. According to this, in a pH close to neutrality, BCECF fluorescence measured as H^+^ activity, or what is the same, H^+^ concentrations using the formula:

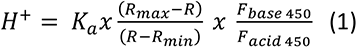

Being K_a_ the acid dissociation constant, R the ratio of the emitted fluorescence by BCECF excited at 490 and 450 (Ratio 490/450), R_max_ is the maximum 490/450 ratio value, obtained in alkaline conditions, R_min_ is the minimum 490/450 ratio in acid conditions. F_base450_/F_acid450_ is the ratio of the fluorescence at 450nm in acid and basic conditions (James-Kracke 1992).

The logarithmic transformation of the equation is:

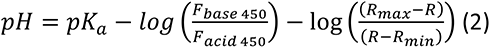

In the isosbestic point F_base450_/F_acid450_ is 1, reason why:

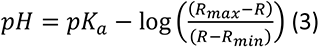

Where R_max_ and R_min_ are obtained from the most alkaline and most acid pH buffers measured.

The values obtained for each culture treated with different buffers were background subtracted and normalized to cell density. The pH calibration curve was obtained by plotting log 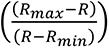 against pH. The resulting equation was used to obtain the pKa. After blank subtraction (yeast without BCECF treatment) and cell density normalization. To obtain pH values, the ratio 490/450 was calculated in each case, and the values obtained were extrapolated from the calibration curve. The data obtained was analyzed with the Microsoft Excel software. Statistical analysis was performed using the OriginPro 2017 program.

### Bioinformatics

A BLAST search against the SwissModel database (Bienert et al. 2017; Waterhouse et al. 2018) using the AtNHX1 protein sequence was conducted to find templates that allow us generate topological and 3D structural models of AtNHX1. The protein sequences of the proteins with better scores identified were obtained from the UniProt database. All pairwise alignments and multiple alignments of proteins sequences performed in this project were conducted using the default parameters of the MUSCLE software (Edgar 2004). Visual 3D model structures were generated using the PyMol software. The evolutionary conservation for AtNHX1 was calculated using the ConSurf server (http://consurf.tau.ac.il/)(Landau et al. 2005). The HMM was generated using the HMMblits tool from the Bioinformatics department of the Max Planck Institute for Developmental Biology, Tübingen (https://toolkit.tuebingen.mpg.de/#/tools/hhblits).

## Acknowledgements

Authors are indebted to Dr. Kees Venema for technical assistance with measurements of vacuolar pH.

---

**Supplemental Figure S1.**
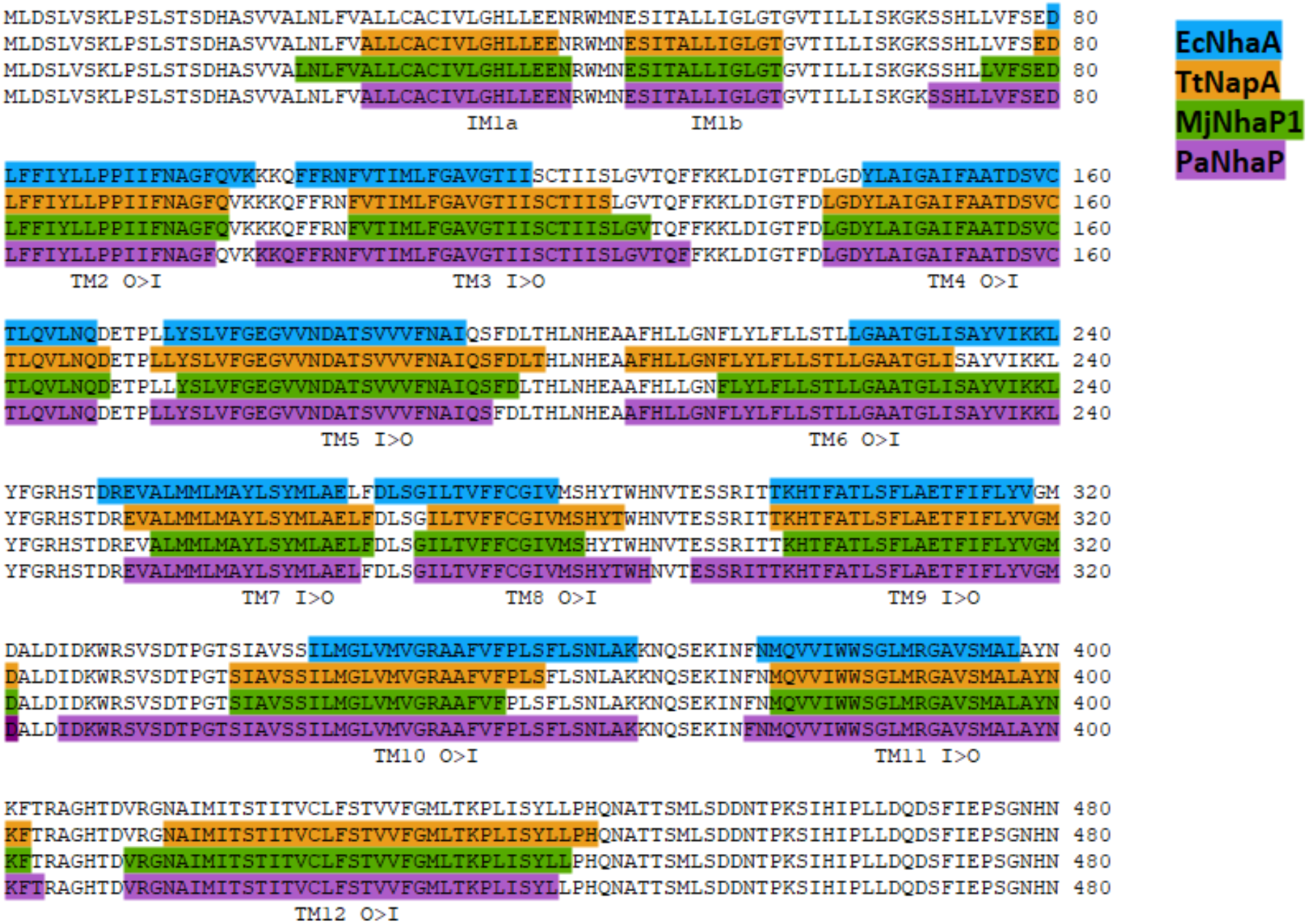
Predicted transmembrane (TM) segments in AtNHX1. Putative TMs are marked according to the alignment with amino acid sequences of crystallized homologous proteins from bacteria and archea. The sequence, repeated four times, corresponds to the AtNHX1 protein on which the corresponding TM segments confirmed in the ternary structure of EcNhaA (blue), TtNapA (orange), MjNhaP1 (green), and PaNhaP (purple) are marked to derive a consensus topology for AtNHX1. The orientation of the twelve TMs predicted in AtNHX1 is marked as outside-to-inside (O>I) or inside-to-outside (I>O).

**Supplemental Figure S2.**
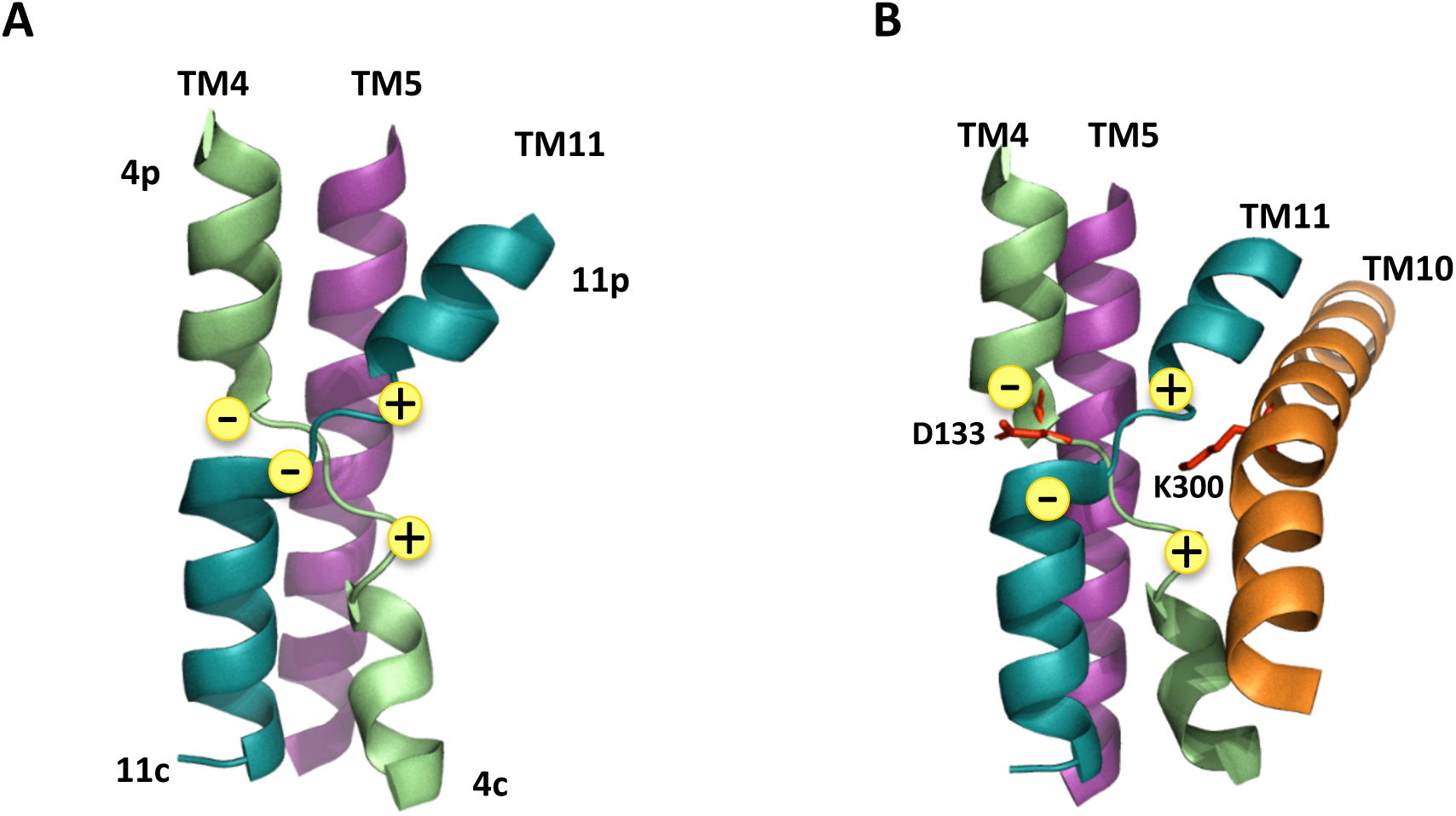
Schematic diagram of the Nha-fold structure. **(A)** TM4 (green) and TM11 (blue) are discontinuous and crossed over the center of the protein, near TM5. The discontinuous helices generate dipoles of opposite charge. **(B)** Asp133 in TM4 and Lys300 in TM10 neutralize the positively and negatively charged helices. Structure is based in the crystal structure of EcNhaA (PDR code 4AU5) (Lee et al., 2014).

**Supplemental Figure S3.**
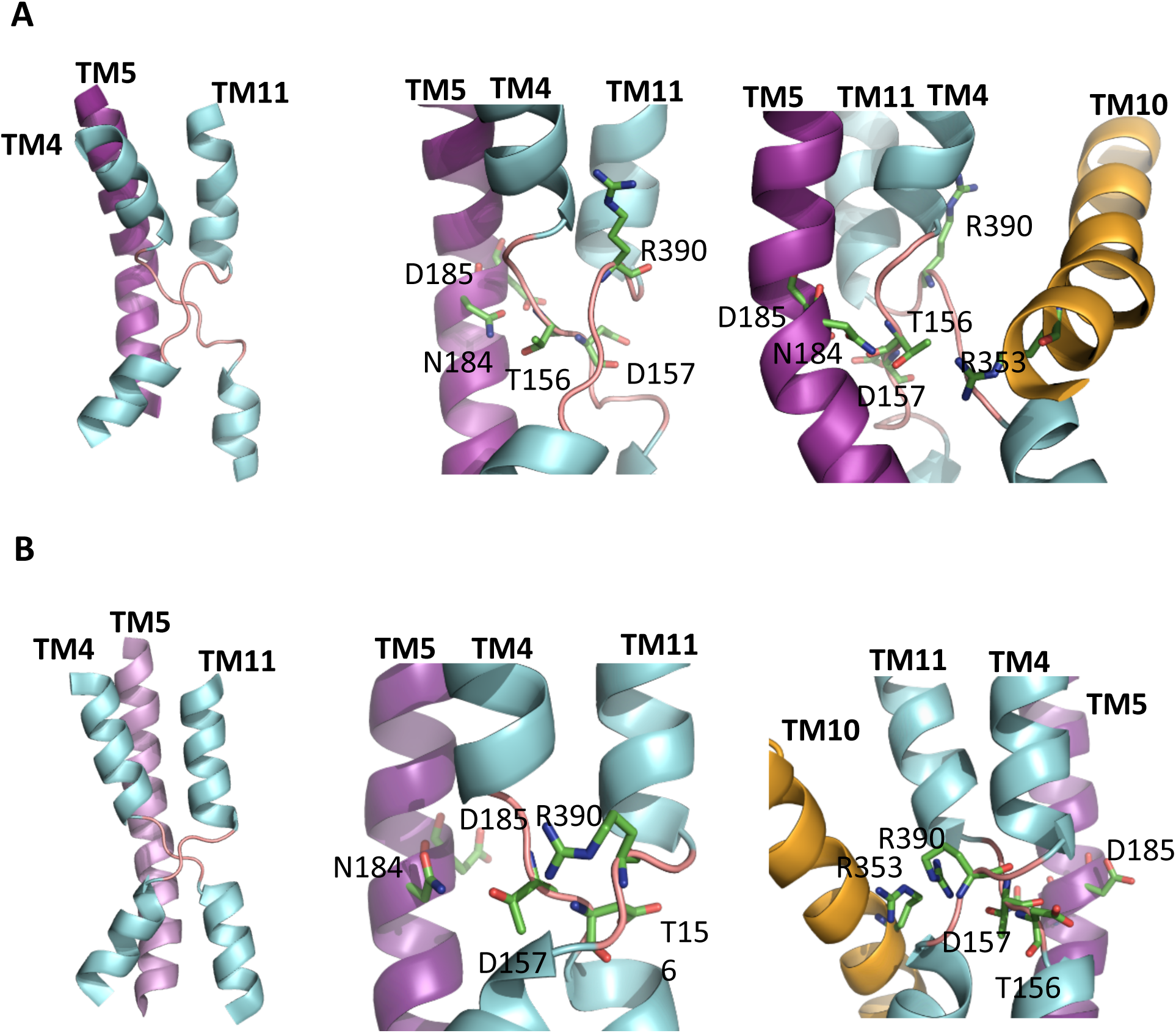
Analysis of the Nha-fold in different models of AtNHX1. Representation of Nha-fold in the active center of the models of AtNHX1 and close-up view of the cross-over of the extended chains of the highly conserved residues T156 and D157 in TM4, N184 and D185 in TM5, and R390 in TM11. Models generated using (**A**) MjNhaP1 (4czb) and **(B)** TtNapA (4bwz) as templates.

**Supplemental Figure S4.**
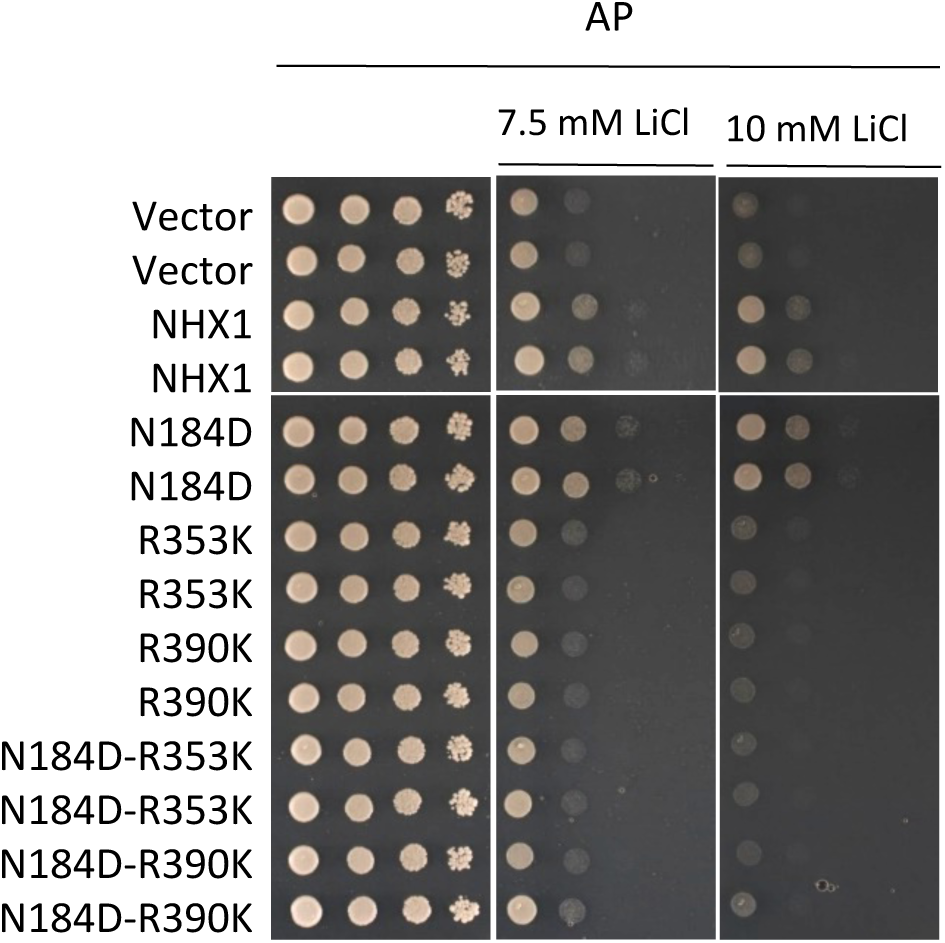
Functional assay of AtNHX1 mutant alleles in amino acids at the ion coordination pocket. The cDNAs of wild-type AtNHX1 and the indicated mutant alleles of the indicated residues were subcloned into the yeast expression vector pDR195 and transformed into the AXT3K (*Δena1-4 Δnha1 Δnhx1*). Overnight cultures were normalized in water to OD_600_ of 0.5. Aliquots (5μL) from normalized cultures and 10-fold serial dilutions were spotted onto AP medium plates. Plates were incubated 2-3 days at 30°C and pictured.

**Supplemental Figure S5.**
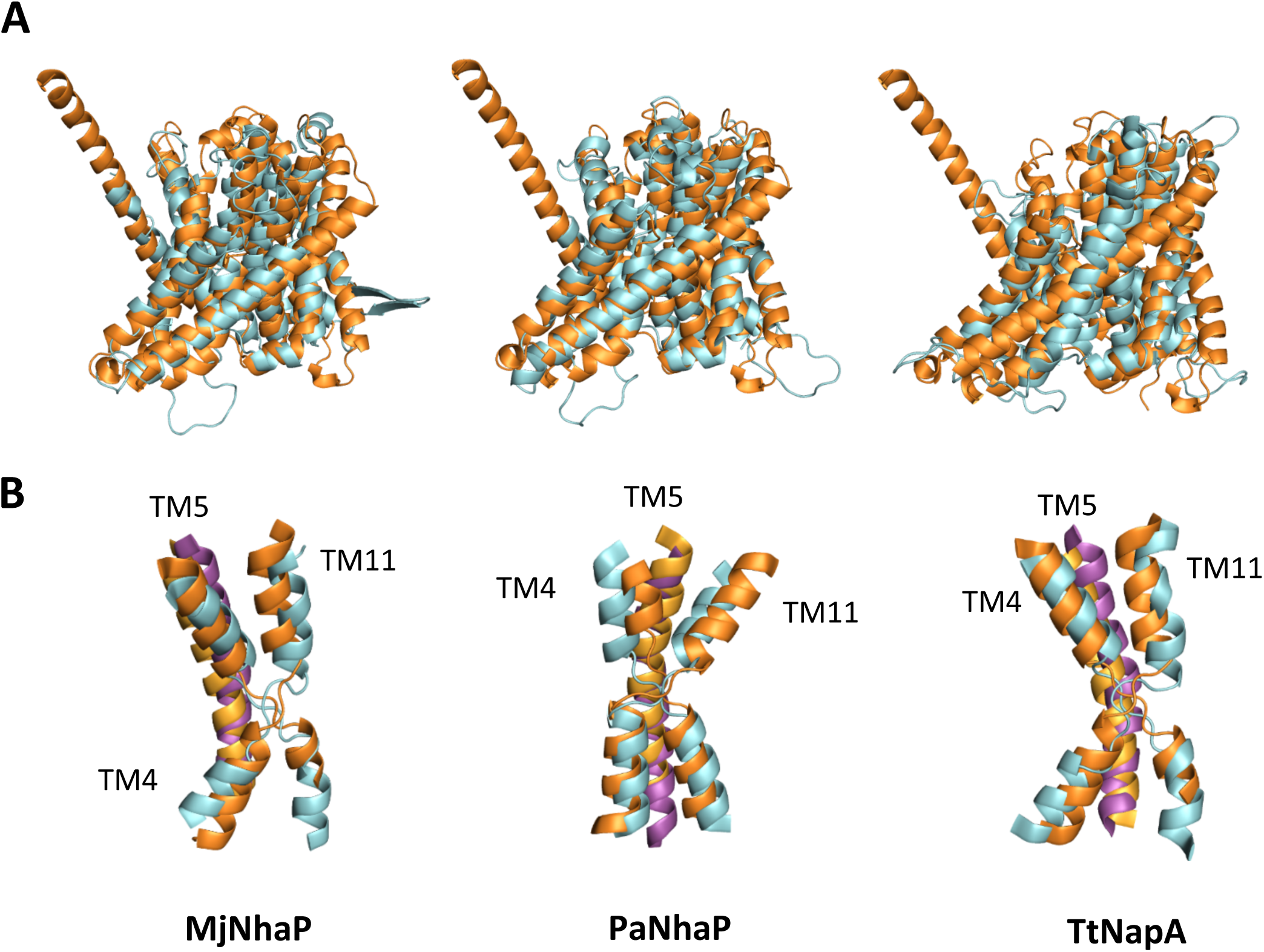
Overlay of the AtNHX1 three-dimensional structures generated by the AlphaFold software (orange) and the different models generated using the proteins in the Swiss-model repository as templates (blue). **(A)** Overlay of the pore domains. **(B)** Overlay of the Nha-fold structures. The names at the bottom indicate template used in each case.

**Supplemental Table S1.**
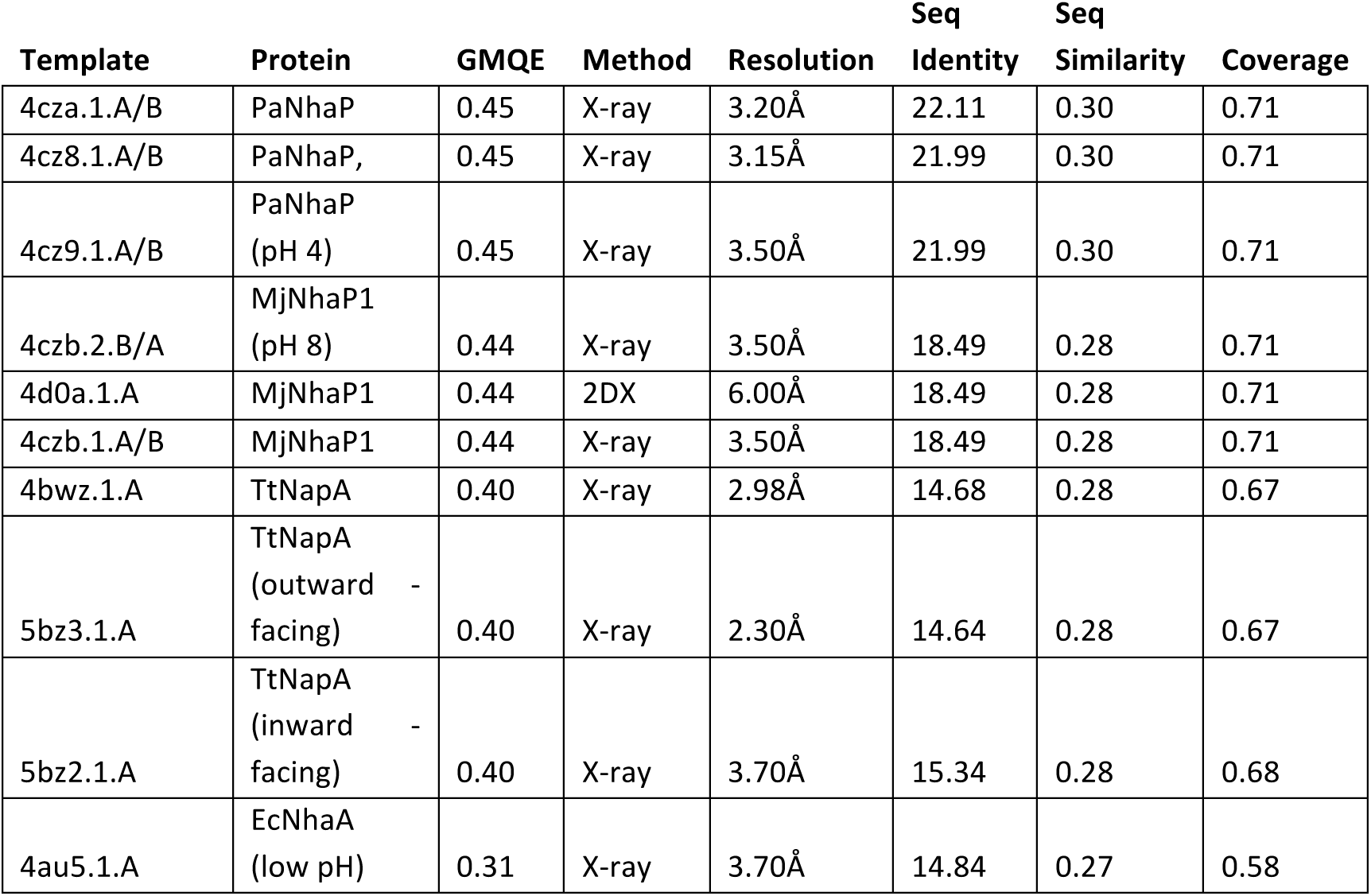
SwissModel best-fitting 3D structures for AtNHX1. Top matches with known Na^+^/H^+^ exchanger structures (templates) are shown with their PDB code. GMQE (Global Model Quality Estimation) is a quality estimation reflecting the expected accuracy of a model built with that alignment of the template and the coverage of the target; Sequence Identity refers to the percentage of amino acid identity between the query and the template; Sequence Similarity is calculated from a normalized BLOSUM62 substitution matrix; Coverage is the proportion of the full- length target protein that is aligned to the template.

| Template | Protein | GMQE | Method | Resolution | Seq<br>Identity | Seq<br>Similarity | Coverage |
| --- | --- | --- | --- | --- | --- | --- | --- |
| 4cza.1.A/B | PaNhaP | 0.45 | X-ray | 3.20Å | 22.11 | 0.30 | 0.71 |
| 4cz8.1.A/B | PaNhaP, | 0.45 | X-ray | 3.15Å | 21.99 | 0.30 | 0.71 |
| 4cz9.1.A/B | PaNhaP<br>(pH 4) | 0.45 | X-ray | 3.50Å | 21.99 | 0.30 | 0.71 |
| 4czb.2.B/A | MjNhaP1<br>(pH 8) | 0.44 | X-ray | 3.50Å | 18.49 | 0.28 | 0.71 |
| 4d0a.1.A | MjNhaP1 | 0.44 | 2DX | 6.00Å | 18.49 | 0.28 | 0.71 |
| 4czb.1.A/B | MjNhaP1 | 0.44 | X-ray | 3.50Å | 18.49 | 0.28 | 0.71 |
| 4bwz.1.A | TtNapA | 0.40 | X-ray | 2.98Å | 14.68 | 0.28 | 0.67 |
| 5bz3.1.A | TtNapA<br>(outward -<br>facing) | 0.40 | X-ray | 2.30Å | 14.64 | 0.28 | 0.67 |
| 5bz2.1.A | TtNapA<br>(inward -<br>facing) | 0.40 | X-ray | 3.70Å | 15.34 | 0.28 | 0.68 |
| 4au5.1.A | EcNhaA<br>(low pH) | 0.31 | X-ray | 3.70Å | 14.84 | 0.27 | 0.58 |

**Supplemental Table S2.**
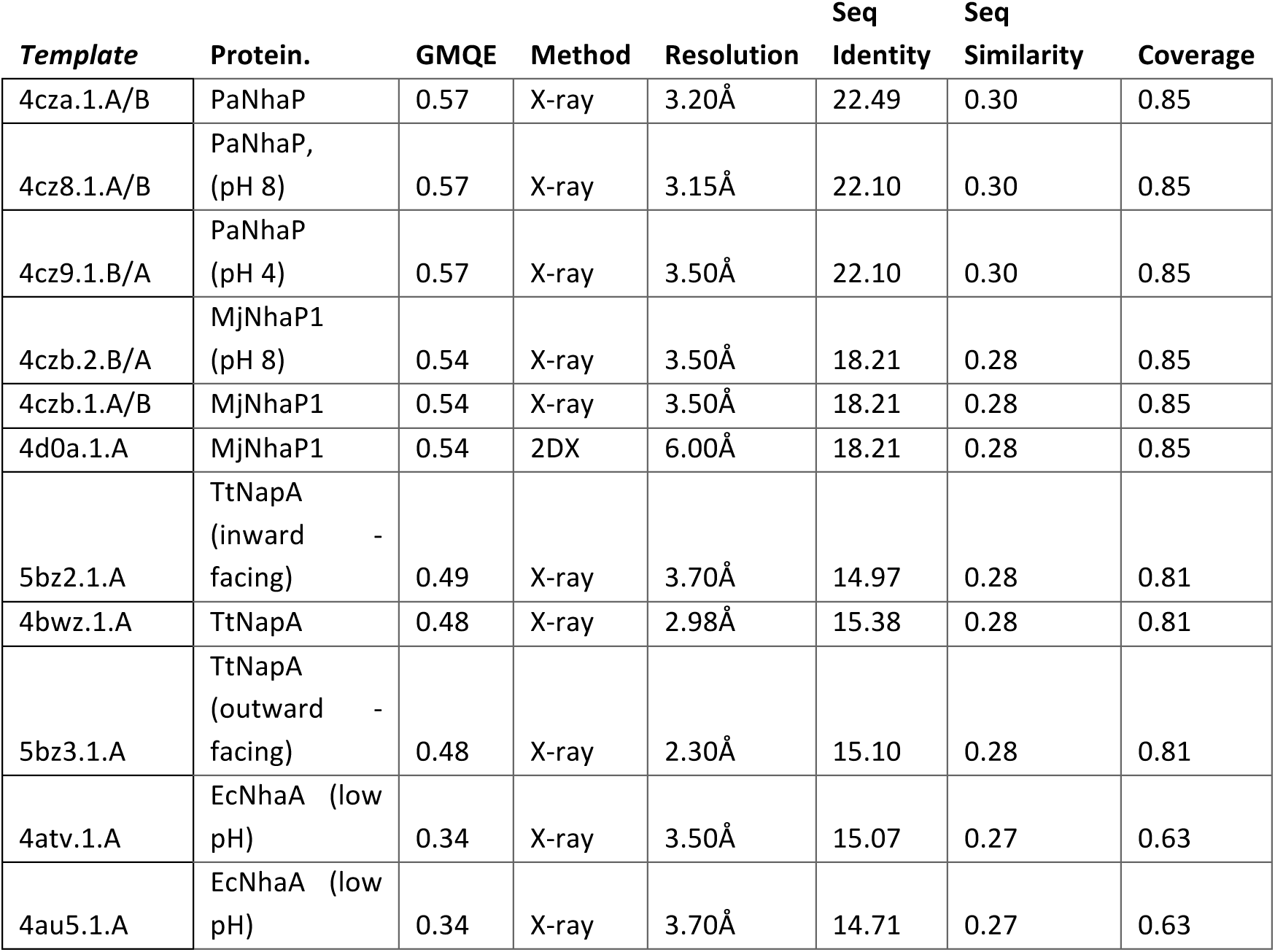
SwissModel best fitting 3D structure for AtNHX1 protein using amino acids 1-435 as query. Top matches with known Na^+^/H^+^ exchanger structures (templates) are shown with their PDB code. GMQE (Global Model Quality Estimation) is a quality estimation that reflects reflecting the expected accuracy of a model built with that alignment and template and the coverage of the target; Sequence Identity refers to the amino acid percentage identity between the query and the template; Sequence Similarity is calculated from a normalized BLOSUM62 substitution matrix; Coverage is the proportion of the full-length target protein that is aligned to the template.

**Supplemental Table S3.**
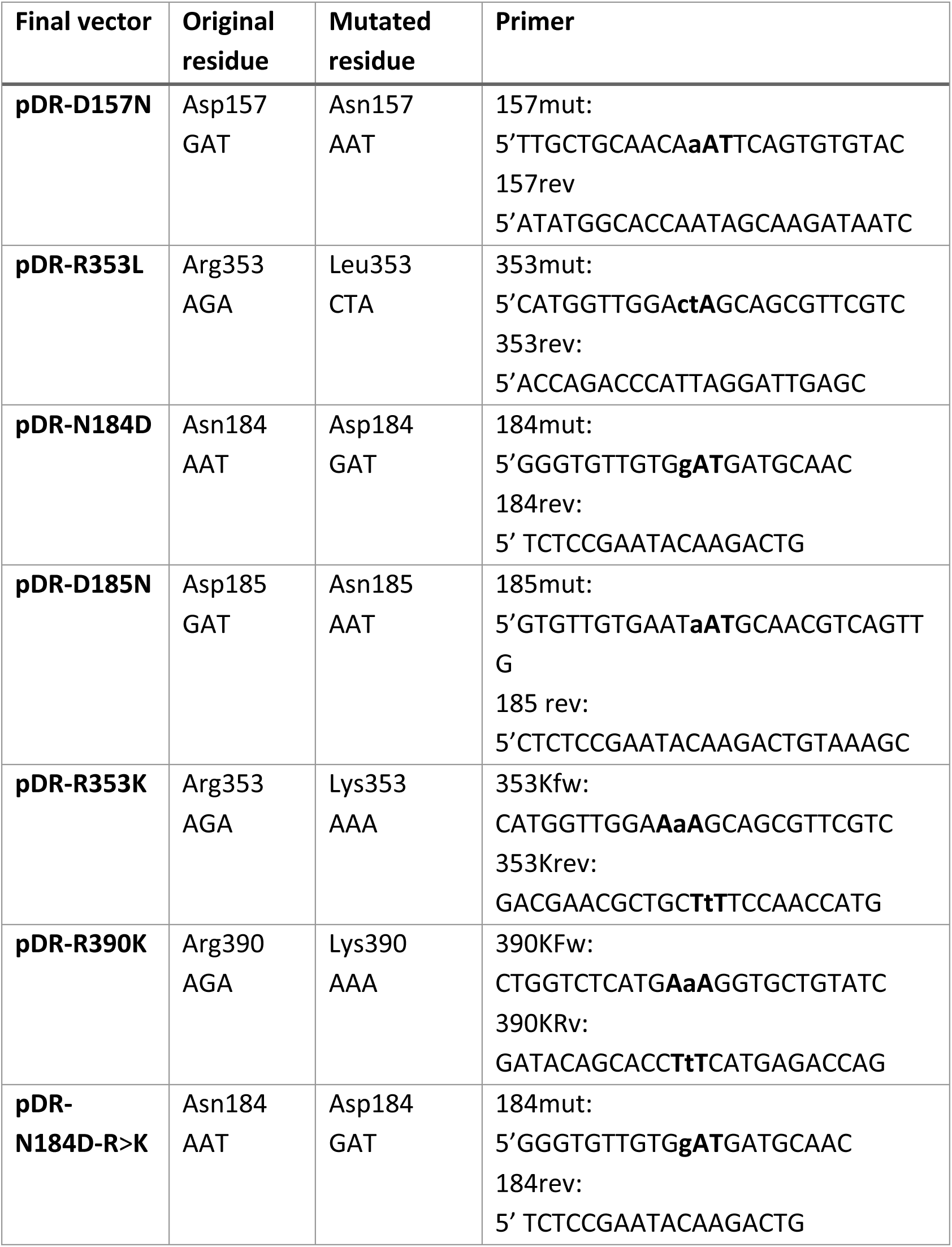
Amino acid residues and primers used for generating NHX1 mutant proteins. Bold triplets indicate the location of the modified codon, and the small letter the change introduced.

## Notes

**Funding:** This work was supported by grant RTI2018-094027-B-I00 from the Spanish Agencia Estatal de Investigación (AEI), of Ministerio de Ciencia, Innovación y Universidades (MCIU), and co-financed by the European Regional Development Fund, to E.O.L. and J.M.P.

### Competing Interest Statement

The authors have declared no competing interest.

